# Sequential Transmission of Task-Relevant Information in Cortical Neuronal Networks

**DOI:** 10.1101/2021.08.31.458395

**Authors:** Nikolas A. Francis, Shoutik Mukherjee, Loren Koçillari, Stefano Panzeri, Behtash Babadi, Patrick O. Kanold

## Abstract

During auditory task performance, cortical processing of task-relevant information enables mammals to recognize sensory input and flexibly select behavioral responses. In mouse auditory cortex, small neuronal networks encode behavioral choice during a pure-tone detection task, but it is poorly understood how neuronal networks encode behavioral choice during a pure-tone discrimination task where tones have to be categorized into targets and non-targets. While the interactions between networked neurons are thought to encode behavioral choice, it remains unclear how patterns of neuronal network activity indicate the transmission of task-relevant information within the network. To this end, we trained mice to behaviorally discriminate target vs. non-target pure-tones while we used *in vivo* 2-photon imaging to record neuronal population activity in primary auditory cortex layer 2/3. We found that during task performance, a specialized subset of neurons transiently encoded intersection information, i.e., sensory information that was used to inform behavioral choice. Granger causality analysis showed that these neurons formed functional networks in which task-relevant information was transmitted sequentially between neurons. Differences in network structure between target and non-target sounds encoded behavioral choice. Correct behavioral choices were associated with shorter timescale communication between neurons. In summary, we find that specialized neuronal populations in auditory cortex form functional networks during auditory task performance whose structures depend on both sensory input and behavioral choice.

## INTRODUCTION

Cortical processing of task-relevant information enables mammals to recognize behaviorally meaningful stimuli while navigating their sensory environment. Performance of an auditory task modulates the neural representations of task-related sounds at the level of single neurons or small populations, already in primary auditory cortex (A1) (Fritz, Shamma et al. 2003, Brosch, Selezneva et al. 2011, David, Fritz et al. 2012, Niwa, Johnson et al. 2013, Rodgers and DeWeese 2014, Kato, Gillet et al. 2015, McGinley, David et al. 2015, Tsunada, Liu et al. 2016, Carcea, Insanally et al. 2017, Kuchibhotla, Gill et al. 2017, Bagur, Averseng et al. 2018, Christison-Lagay and Cohen 2018, Francis, Elgueda et al. 2018, Francis, Winkowski et al. 2018, Schwartz and David 2018, Guo, Weems et al. 2019, Insanally, Carcea et al. 2019, Yin, Strait et al. 2020). We recently showed that performing a pure-tone detection task increases neuronal responses to target tones in A1 layer 2/3 (L2/3) and changes the functional network connectivity in L2/3 by forming small strongly linked neuronal networks that encode behavioral choice (Francis, Winkowski et al. 2018). However, natural auditory scenes typically include both target and non-target sounds that require discrimination from each other. The effect of this discrimination on the functional networking of neurons and how such target vs. non-target information is propagated through the network are poorly understood.

Given the diversity of neuronal connectivity and stimulus selectivity in A1 L2/3 (Atzori, Lei et al. 2001, Yang, DeWeese et al. 2008, Sadagopan and Wang 2009, Sakata and Harris 2009, Atencio and Schreiner 2010, Bandyopadhyay, Shamma et al. 2010, Oviedo, Bureau et al. 2010, Rothschild, Nelken et al. 2010, Winkowski and Kanold 2013, Kanold, Nelken et al. 2014, Maor, Shalev et al. 2016, Meng, Winkowski et al. 2017), we hypothesized that there may exist specialized neurons in A1 L2/3 that represent varying amounts of sensory or choice information. Moreover, we hypothesized that a subset of these neurons carry sensory information used to inform behavioral choice and form functionally connected networks whose interactions encode behavioral choices during task performance. To investigate our hypotheses, we trained mice to behaviorally discriminate target vs non-target pure-tones while we recorded neuronal activity in A1 L2/3 using 2-photon (2P) Ca^2+^ imaging. We then quantified how much stimulus information (*SI*), behavioral choice information (*CI*), and intersection information (*II*), i.e., sensory information that is used to inform behavioral choice, was carried by individual neurons (Panzeri, Harvey et al. 2017, Runyan, Piasini et al. 2017). We also used Granger causality (GC) analysis to study how these neurons were organized into functional networks (Kaminski, Ding et al. 2001, Bressler and Seth 2011, Kim, Putrino et al. 2011, Quinn, Kiyavash et al. 2015, Seth, Barrett et al. 2015, Francis, Winkowski et al. 2018, Sheikhattar, Miran et al. 2018). Here, we extended GC analysis to study both the functional networks and the timescales of information processing in A1 functional networks.

We found that task performance modulated neuronal response amplitudes, network structures, and network information transmission in A1 L2/3. Neuronal populations that encoded *II* exhibited sparse network connections and encoded task-relevant information at different peak times that tiled the duration of a trial. Networked neurons encoding *II* shared redundant sound information relevant for behavioral choice. Together, our results describe how neurons in A1 L2/3 that integrate sensory and behavioral information during auditory task performance form functional connections that sequentially transmit task-related information between networked neurons.

## RESULTS

To study how task-relevant information is transmitted within neuronal networks, we trained 9 transgenic CBA x Thy1-GCaMP6s F1 mice (Frisina, Singh et al. 2011, c17Dana, Chen et al. 2014) to perform a pure-tone frequency discrimination task while we imaged neuronal responses in A1 L2/3 using *in vivo* 2P Ca^2+^ imaging (Fig. 1).

**Figure 1.**
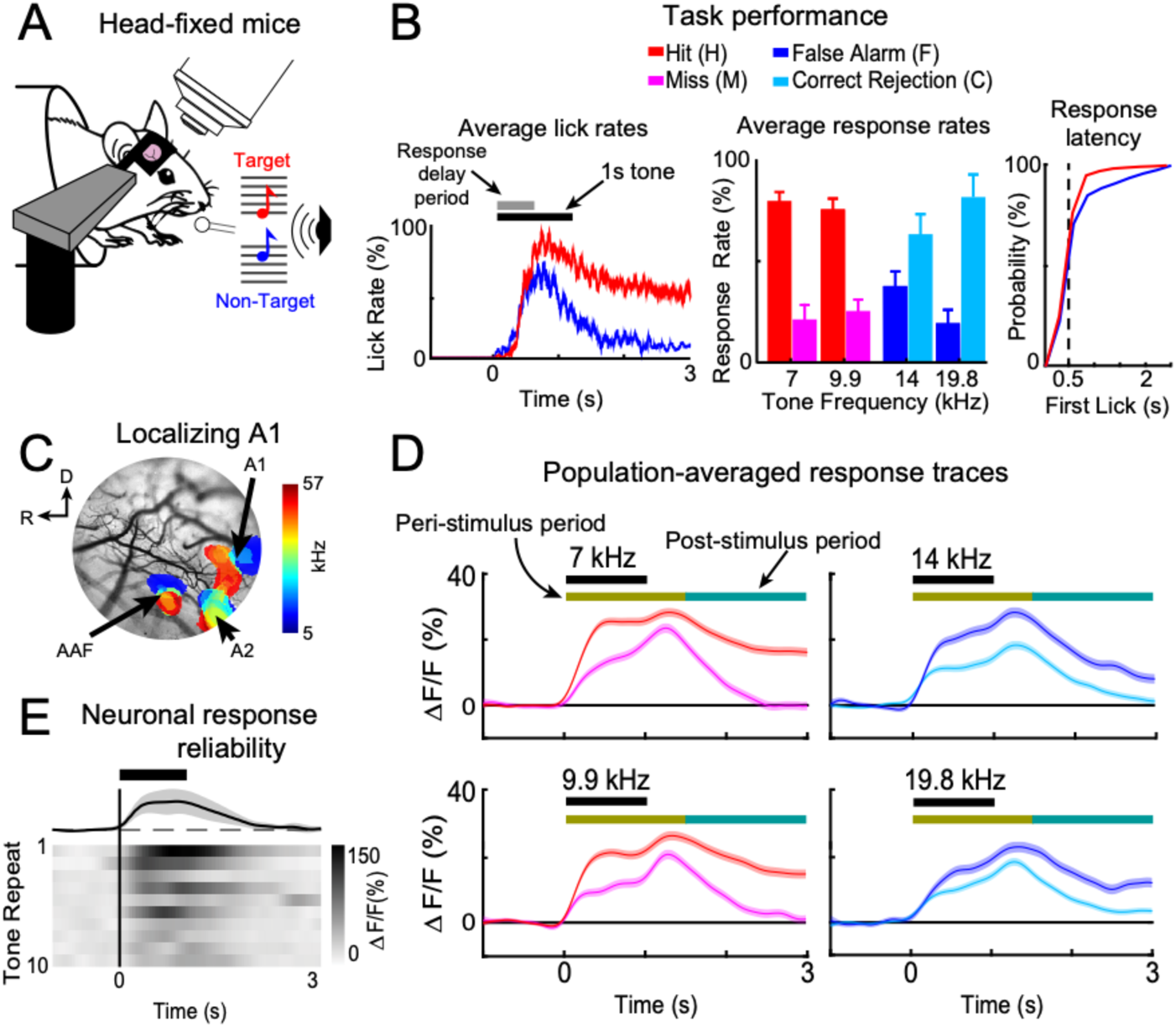
2-photon (2P) imaging in awake-behaving mice shows that neural responses are modulated by behavioral choice. **A.** Head-fixed mice were trained to discriminate low-frequency target tones (red) vs high-frequency non-target tones (blue). **B.** The left panel shows the average lick rates within a trial during task performance. Horizontal black and gray bars show the tone presentation and behavioral response-delay periods, respectively. The red trace shows the lick rate for hits (H). The blue trace shows the lick rate for false alarms (F). The middle panel shows the average behavioral-choice rates, i.e., Hit (red), Miss (pink), False Alarm (blue), and Correct Rejection (cyan), for each presented tone frequency. Error bars show two standard errors of the mean (SEMs; n=34 experiments). The right panel shows the cumulative distribution functions across experiments for hit (red) and false alarm (blue) behavioral response latencies. **C.** Primary auditory cortex (A1) was localized within a craniotomy by using wide-field imaging to visualize tonotopy in auditory cortex. **D.** Average neuronal population response traces in A1 layer 2/3 (L2/3) (N=2692 neurons) color-coded for behavioral choice as in panel B. Each trace shows the response to the pure-tone frequency shown above the black horizontal bar that indicates when the tone was presented. Shading shows 2 SEMs. The horizontal colored bars show the peri-, and post-stimulus windows, respectively, used for later analyses. **E.** Neurons in A1 L2/3 responded transiently, with jittered amplitude and timing in response to repeated identical pure-tones.

### Head-fixed mice learned to perform an auditory frequency discrimination task

Head-fixed mice were trained to lick a waterspout in response to hearing a low-frequency target tone (Fig. 1A; 7 or 9.9 kHz, red), and to avoid licking the waterspout after hearing a high-frequency non-target tone (14 or 19.8 kHz, blue). The four frequencies were randomly interleaved across trials. Water-restricted mice were trained to delay behavioral responses for 0.5 s after the onset of a target tone to receive a water droplet. This waiting period enabled us to assess neural responses before the animal indicated its decision by licking, and thus minimizing task-related motor signals in A1 (Francis, Winkowski et al. 2018).

Figure 1B shows that the mice learned to behaviorally discriminate targets vs. non-targets. Each trial’s behavioral response was categorized into four possible behavioral choices: hit (H: licking after target onset), miss (M: no licking after a target), false alarm (F: licking after non-target onset), or correct rejection (C: no licking in response to a non-target). The right panel of figure 1B shows the cumulative distribution functions for H and F behavioral response times (i.e., the time of the first lick in a trial) in each experiment. The average H and F response latencies relative to stimulus onset were 0.64s ± 0.02s and 0.75s ± 0.04s, respectively. 74% of H trials had behavioral response latencies greater than 0.5s after stimulus onset. Thus, the mice usually delayed behavioral responses until 0.5s after stimulus presentation during target trials to receive a reward. On average across 34 experiments, both the overall hit rate (78.8% +± 5.1%) and the rewarded hit rate (H = 58.3% ± 6.6%) were significantly greater than the false alarm rate (F = 27.1% ± 7.3%; p<0.001, t-test). Similarly, the correct rejection rate (74.3% ± 6.9%) was significantly higher (p<0.001, t-test) than both the F and M rate (20.8% ± 5.2%). These data show that the mice were able to discriminate between target vs. non-target sounds.

### Decision-making modulated neuronal response amplitude in A1 L2/3

To characterize neural responses during behavior, we imaged Ca^2+^-dependent fluorescence in auditory cortex. To localize 2P imaging fields for each experiment to A1, we first mapped the frequency organization of auditory cortex (i.e., tonotopy) in each mouse using widefield (WF) imaging (Fig. 1C) (Francis, Winkowski et al. 2018, Liu, Whiteway et al. 2019).

To identify cellular responses in A1 L2/3, we performed 2P imaging (Fig. 1D,E) at a depth of 150-250 µm from the cortical surface in each mouse (34 experiments, 9 mice, N = 2792 neurons). We observed fluorescence (ΔF/F) responses to tones at all 4 task frequencies (7, 9.9, 14, and 19.8 kHz), with response dynamics typical of GCaMP6s fluorescence (Chen, Wardill et al. 2013, Dana, Chen et al. 2014). The neural traces showed a complex pattern of task-dependent changes in response amplitude. Traces typically began to rise quickly during the first few hundred milliseconds of the tone onset, and the post-stimulus decay-rate varied across conditions. In general, fluorescence responses to non-target frequencies (14 and 19.8 kHz) were smaller in amplitude than responses to target frequencies (7 and 9.9 kHz) (p<0.001). In addition, trials without behavioral responses, i.e., misses and correct rejections, had lower average response amplitudes than trials with behavioral responses, i.e., hits and false alarms. Thus, the amplitude of pure-tone responsiveness in A1 during task performance was strongly modulated not only by acoustic stimulation, but also by behavioral choice.

### Task-relevant information is transiently encoded by individual neurons in A1 L2/3

We hypothesized that different neurons in A1 might represent varying amounts of sensory or choice information. To quantify the task-relevant information carried by each neuron, we used information theory at the single-trial level (Shannon 1948, Quian Quiroga and Panzeri 2009). For each neuron we quantified how much information was present about the sound stimulus (*SI;* i.e., target vs non-target tone; Fig. 2A, top) and about the behavioral choice (*CI*; i.e., lick vs no-lick; Fig. 2A, bottom). We also computed intersection information, (*II*; Fig. 2A, middle) (Panzeri, Harvey et al. 2017, Pica, Piasini et al. 2017), which quantifies how much of the sensory information encoded by the neurons is used to inform behavioral choices, and is thus a direct measure of task-relevant information.

**Figure 2.**
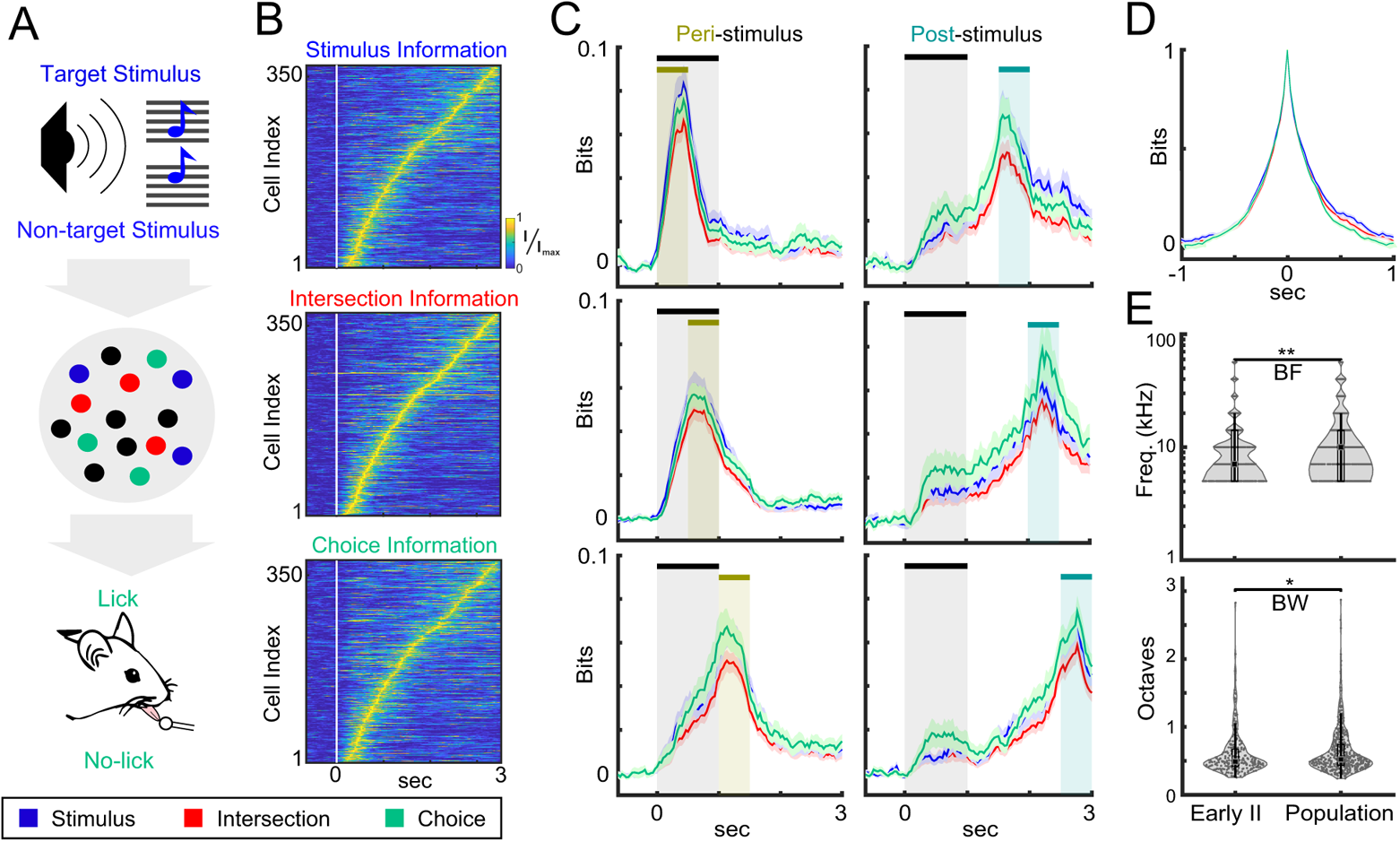
A1 L2/3 neurons transiently carried stimulus, choice and intersection information. **A.** Schematic representation of the information framework shows the two stages of information processing for auditory task performance, namely stimulus encoding and behavioral read out, based on the neural activity of A1 L2/3 neurons. Blue, green and red circles respectively represent neurons with stimulus information (*SI*) only, choice information (*CI*) only, and intersection information (*II*). Intersection information accounts for the part of sensory and choice information used to perform the task. **B.** Information time-courses for *SI*, *CI*, and *II* were normalized to the peak of each neuron’s information (*I_max_*) and sorted by peak time of *II*. **C.** Time-course of *SI, CI*, and *II* averaged over neurons showed similar patterns. We quantified the *SI, CI,* and *II* in six separate stages of the behavioral task, which account for the *peri-stimulus* (0-1.5 s) and the *post-stimulus* intervals (1.5-3 s) shown by the shaded regions. Error bars show one standard error of the mean (SEM; N=#neurons with *II* peaks within the stage). **D**. Transiency of *SI, CI,* and *II*, computed by aligning information peaks across neurons and then analyzing the peak-aligned traces within ± 1 s from the peak. **E**. Violin plots of the estimated best frequency (BF) (top) and tuning bandwidth (BW) (bottom) of neurons with early *II* vs. other neurons in the observed population. Early *II* neurons have significantly lower BF (p < 0.01, Wilcoxon rank sum test) compared to the other neurons in the population, consistent with the low frequency target stimuli. They also have significantly narrower selectivity bandwidth (p < 0.05, Wilcoxon rank sum test), suggesting sharper tuning to the target frequency.

We identified the times at which neurons carried *SI, CI,* and *II*, by first computing deconvolved spike rates, obtained with a sliding window approach across the entire duration of the trial (see Methods). We inferred the relative spiking activity from the fluorescence traces with a first-order autoregressive model using the CaImAn algorithm (Vogelstein, Packer et al. 2010, Giovannucci, Friedrich et al. 2019). Figure 2B shows the peak-normalized information time-courses for each neuron, sorted by the peak-latency of the respective information (*SI, CI* or *II*) across neurons. We found that 1183 neurons (42.37% of the 2792 recorded neurons) had significant *SI* or *CI* (permutation test, p < 0.1, corrected for comparisons across multiple time windows, see Methods) and thus carried information about the stimulus or the behavioral choice. Of the 1183 neurons carrying either stimulus or choice information, 708 neurons did not carry significant intersection information (permutation test, p > 0.1, corrected for comparisons across multiple time windows), meaning that these neurons either had stimulus information that did not inform choice (e.g. because that cell’s stimulus signal was not causal to formation of the choice) or choice information not related to the stimulus (e.g. internal choice bias). The remaining 475 (40.15%) carried significant *II* (permutation test, p < 0.1, corrected for comparisons across multiple time windows), and thus integrate both sensory and behavioral information that is directly relevant for the decision-making task (Panzeri, Harvey et al. 2017). Moreover, qualitative inspection showed that neurons carried *SI*, *CI*, or *II* transiently and sequentially, tiling the trial duration.

We restricted our further analyses to the sessions (12 out of 34 sessions) with at least 20 *II* neurons due to our interest in subsequent network analyses for which, given the number of experimental trials, at most 20 neurons could be analyzed with statistical confidence. This left us with 375/475 significant *II* neurons for our subsequent analyses (see Methods). The *SI, CI,* and *II* time-courses of high *II* neurons showed similar population-averaged time patterns and for neurons carrying all three types of information the peak of information of each information type was similar (Fig. 2C), indicating that each cell had a specific peak time point at which most or all its task-relevant information was expressed. Neurons that carried either significant *SI* or *CI* but not *II*, showed similar trends in their *SI and CI* time-courses (Figure 2 – Figure Supplement 1). To identify how much of the stimulus information is used for informing behavioral choice, we computed the ratio between *II* and *SI*. Of the neurons with significant *II*, the average *II/SI* ratio was high (>70%) throughout a trial, meaning that most of the *SI* in these single neurons in A1 L2/3 was used for the task. These results indicate that a considerable fraction of A1 neurons transiently carry significant stimulus information that is used to inform behavioral choice.

Neurons had transiently high *SI*, *CI*, or *II*, but across the population different neurons carried *II* at different times during the trial. To shed light on the temporal dependencies of information carried by individual neurons across the phases of the trial, we defined two types of neurons based on their *II* peak latencies. We labelled neurons that peaked in the first 1.5 seconds after stimulus onset as *peri-stimulus informative*, and the remainder as *post-stimulus informative* (Fig. 2C). We subdivided the *peri-stimulus informative* neurons into three sequential task-related periods within a trial: (1) the 500 ms waiting period just after tone onset, (2) the 500 ms interval after the waiting period, and (3) the 500 ms after tone offset (Fig. 2C, left column). We found that 52/375, 85/375, and 60/375 neurons had *II* that peaked in the first, second, and third *peri-stimuli* periods, respectively (Fig. 2C, left column). Thus, 197/375 individual neurons encoded *II* during the *peri-stimulus* interval. 133/375 neurons had *II* that peaked in the *post-stimulus* period (1.5-3 s) with 45/375, 40/375, and 48/375 neurons in the first, second, and third *post-stimulus* periods, respectively (Fig. 2C, right column). The remainder 45/375 neurons peaked after 3 s. Although the values of *SI*, *CI* and *II* remained comparable throughout the trial, neurons with peaks in the pure tone presentation period carried slightly more *SI* than *CI* (Fig. 2C, left column, blue vs. green traces), and neurons with later response times carried slightly higher *CI* than *SI* (Fig. 2C, right column, green vs. blue traces).

Qualitative inspection of the *SI*, *CI*, or *II* time-courses suggested that neurons transiently carried information. To quantify the transiency of information coding, we aligned information peaks across neurons and then analyzing the peak-aligned traces within ± 1 s from the peak (Fig. 2D).

The peak-aligned traces admit an exponential fit with a time constant 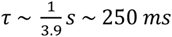. Thus, individual neurons transiently carried *SI, CI, II* for an effective duration of ~250 *ms*.

Given that *II* neurons carried *SI* we compared the tuning of *II* neurons (Fig. 2E). We found that the best frequencies (BFs; the sound frequency values eliciting the highest response during passive pure-tone presentation) of *II* neurons were lower (p<0.01, Wilcoxon rank sum test) than the average BF of the general population of all imaged neurons. *II* neurons also had narrower bandwidth (BW) (p<0.05, Wilcoxon rank sum test) than the general population. Thus, we found that *II* neurons tended to be sharply tuned and had BFs that were biased toward the low target frequencies in our auditory task.

Together, our results show that a subset of neurons in A1 L2/3 encoded task-relevant information. Importantly, task-relevant information was transiently encoded by individual neurons, yet sustained throughout the trial by sequential encoding across the population.

### Neurons with intersection information form sparse timescale-invariant networks

So far, we have shown that a large fraction of individual neurons in A1 L2/3 encode *II*, i.e. sensory information that is relevant for behavioral choice. Since individual neurons had low information content and only transiently encoded *II* (Fig. 2B,C), we hypothesized that *II* neurons might form functional networks to more robustly encode task-relevant information and that these networks persist over time. To determine if neurons organized into functional networks, we used Granger causality (GC) analysis. We previously used GC analysis to identify small networks of interacting neurons whose network structure depended on behavioral choice (Francis, Winkowski et al. 2018), but we did not study how neuronal network structure might vary with respect to integration timescales. This is important in our go/no-go task design, wherein the mice had to delay behavioral responses until 0.5s after stimulus onset. Moreover, our finding that *II* was transiently encoded by individual neurons, but sustained across time by the population, necessitates the examination of relevant timescales of interactions between *II* neurons. Hence, we extended our previous GC analysis by studying functional network connectivity across multiple timescales. GC analysis uses multivariate statistics to infer significant causal influences within a population of neurons by testing if the recent history of a neuron can improve the prediction of another neuron’s activity. The duration of the recent history over which interactions are quantified, referred to as the ‘integration window’, is a hyperparameter of GC analysis, whose value, d, sets the longest interaction delay considered (Fig. 3A, left schematic). Short (S; d=233 ms) integration windows quantify interactions that are more likely to reflect local neuronal interactions. Long (L; d=1033 ms) integration windows would additionally capture the local effects of potentially slower and indirectly mediated interactions that may involve distant neurons, perhaps reflecting non-sensory task-related interaction such as error signaling or other forms of top-down processing. The specific window lengths we used were integer multiples of the imaging frame rate. Importantly, the S-timescale interactions are a subset of the L-timescale interactions (see Methods). For each experiment (N=12), we performed GC analysis on the 20 neurons with the lowest *II*-peak latencies to identify the contribution of neurons whose activity carried task-relevant information during stimulus presentation. Our GC analysis used 20 neurons per experiment as this was the largest network size for which we were confident that there was no overfitting the data during sparse vector autoregressive modelling, given the number of experimental trials (see Methods). GC networks were estimated individually for each discrimination task behavioral choice category (H, M, C, and F). This importantly contrasts previous work (Francis, Winkowski et al. 2018) in which we could only analyze networks corresponding to *detection* task behavioral choice categories, H and M.

**Figure 3.**
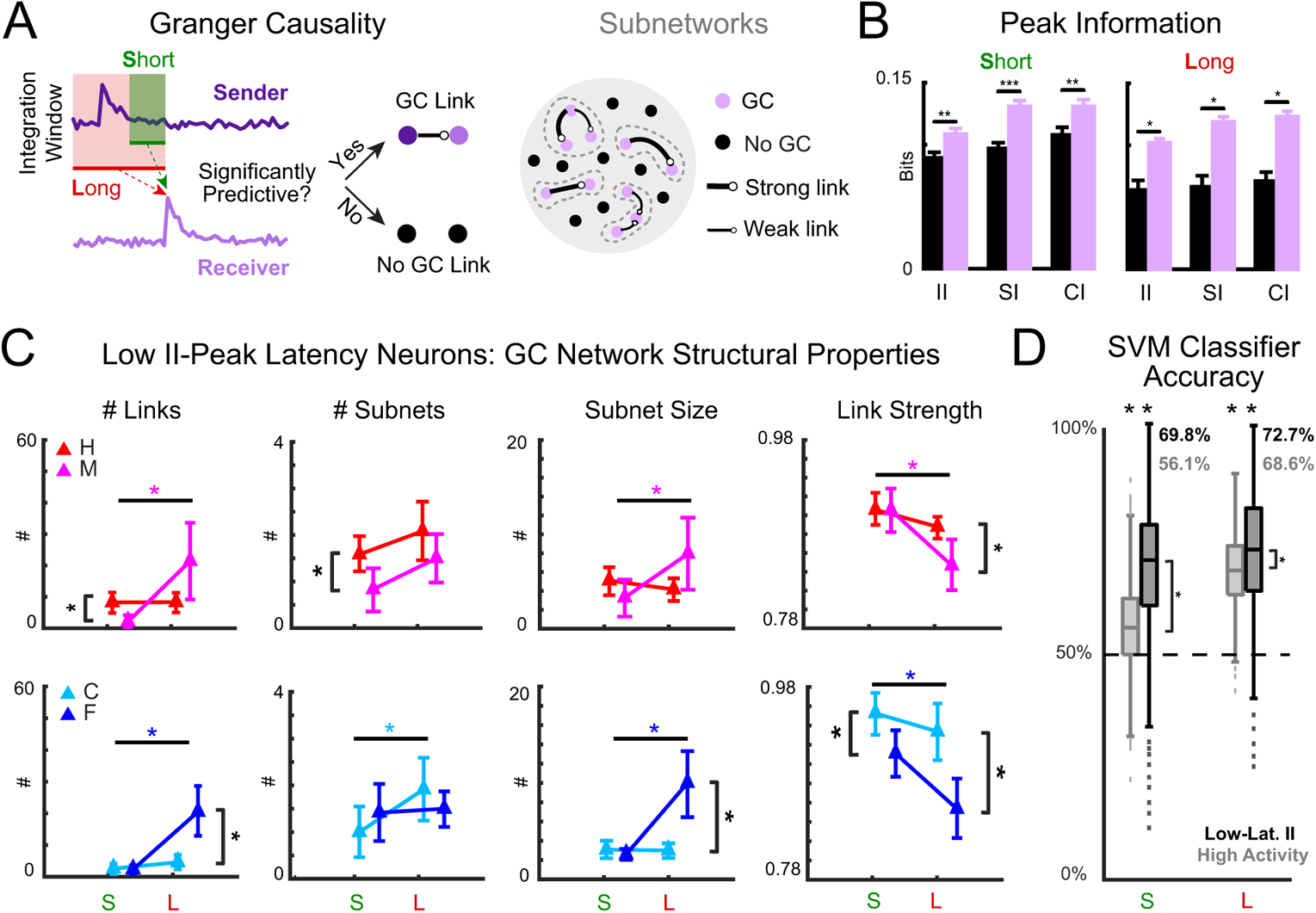
Behavioral choice is encoded in network structure of low *II*-peak latency neurons. **A.** Functional networks of short (S)- and long (L)-timescale interactions amongst low *II*-peak latency neurons were estimated using Granger Causality (GC) analysis for each behavioral choice: Hit (H), Miss (M), Correct Rejection (C), and False Alarm (F). Disjoint sets of interlinked neurons constituted subnetworks (dashed grey boundaries). **B.** GC-linked neurons, for both S and L timescales, had more information than GC-unlinked neurons (*p<0.05; **p<0.01; ***p<0.001). **C.** Four GC network statistics were analyzed: number of links, number of subnetworks, size of subnetworks, and statistical strength of links. For S v. L timescales, correct decision (H/C) networks had robust number of links, subnetwork sizes, and link strengths; incorrect decision (M/F) networks had greater number of links and subnetwork size, and weaker links in L-timescale networks. S-timescale H/C networks had at least as many links as M/F networks, while L-timescale H/C networks had fewer, stronger links. See also Table 1. **D.** Network statistics were used to train an SVM to classify behavioral responses into correct or incorrect decisions. Across timescale and selection of neurons, decisions were predicted significantly better than chance (p<0.001). S-timescale network structure of low *II*-peak latency neurons was better decoded than from highly responsive neurons (p<0.001). In general, L-timescale network structures had high decoding accuracy, but low *II*-peak latency neuronal networks were better decoded than highly responsive neurons (p<0.001). Error bars show 2 SEMs. The **\*** indicates differences statistically significant at p<0.05. Wilcoxon’s signed rank test was used in **C**, and a two-sample t-test in **D**.

**Table 1.** Statistical comparisons of GC network structure across short (S) and long (L) timescales, and behavioral choice categories — hit (H), miss (M), correct-rejection (C), and false-alarm (F) – using Wilcoxon’s signed rank test (p<0.05). See also Fig. 3C.

|  | # Links |  |  | # Subnets |  |  | Subnet Size |  |  | Link Strength |  |  |
| --- | --- | --- | --- | --- | --- | --- | --- | --- | --- | --- | --- | --- |
|  | S | L | p-value | S | L | p-value | S | L | p-value | S | L | p-value |
| H | 8.25 ± 1.61 | 8.33 ± 1.49 | 0.97 | 1.58 ± 0.19 | 2.08 ± 0.31 | 0.19 | 5.11 ± 0.75 | 4.16 ± 0.61 | 0.333 | 0.91 ± 0.009 | 0.89 ± 0.006 | 0.111 |
| M | 2.08 ± 1.05 | 21.42 ± 6.10 | **0.009** | 0.83 ± 0.24 | 1.50 ± 0.26 | 0.074 | 3.30 ± 0.99 | 7.94 ± 1.91 | **0.041** | 0.91 ± 0.012 | 0.85 ± 0.014 | **0.014** |
| p-value | **0.005** | 0.059 |  | **0.024** | 0.167 |  | 0.161 | 0.073 |  | 0.975 | **0.02** |  |
| C | 2.58 ± 0.91 | 4.58 ± 2.32 | 0.189 | 1.00 ± 0.28 | 1.92 ± 0.34 | **0.047** | 3.08 ± 0.42 | 3.00 ± 0.43 | 0.59 | 0.95 ± 0.011 | 0.93 ± 0.015 | 0.369 |
| F | 2.33 ± 0.73 | 20.5 ± 3.90 | **<0.001** | 1.42 ± 0.31 | 1.50 ± 0.19 | 0.824 | 2.59 ± 0.32 | 10.0 ± 1.72 | **<0.001** | 0.91 ± 0.012 | 0.85 ± 0.016 | **0.013** |
| p-value | 0.832 | **0.002** |  | 0.33 | 0.298 |  | 0.357 | **<0.001** |  | **0.049** | **0.002** |  |

We found that across all trials GC networks were sparsely connected: only 1% of possible links connecting 21.98 % of the selected *II* neurons were detected in S-timescale networks, while 3.61% were detected in L-timescale networks connecting 51.67 % of the selected *II* neurons. Unlike simpler measures such as Pearson correlation, GC is a directed measure of communication, which can distinguish senders from receivers (Figure 3A). This allowed us to investigate the proportion of senders and receivers among the neurons in the network. For the S-timescale networks, 10.10% of neurons were senders, 9.06% were receivers, and 2.81% were GC-linked neurons that had net degree of zero. For the L-timescale networks, however, 24.58% of the neurons were senders, 19.79% were receivers and 7.29% had a net degree of zero. This indicates that an additional 29.69% of the selected *II* neurons were recruited over longer integration window.

Speculating that the information content of GC-linked neurons differed from GC-unlinked neurons, we compared *SI, CI,* and *II* at the time when neurons peaked in *II* and found that they were higher in GC-linked neurons than in GC-unlinked neurons in both S-timescale networks and L-timescale networks (Figure 3B). These results suggest that GC-linked neurons form networks carrying signals of greater relevance for performing the auditory discrimination task.

To characterize how the structure of the GC networks formed by neurons carrying task-relevant information depends on the timescale of interactions and on behavioral choice, we analyzed 4 GC network statistics separately for H, M, C, and F trials: number of links, number of subnetworks (isolated subsets of neurons), subnetwork size (number of member neurons), and statistical strength of links (by Youden’s *J*-statistic) (Francis, Winkowski et al. 2018) (shown from left to right in Fig. 3C; see also Table 1). For both M and F networks, the number of links and the size of subnetworks were greater for L- than S-timescale networks, while link strength was less for L- than S-timescale networks. In contrast, we found no differences for L- vs. S-timescale networks in H or C trials for the number of links, size of subnetworks, and link strengths. In C trials, number of subnetworks increased with integration window length. Together, our results show that incorrect decision (e.g. M and F) L-timescale networks are larger but connected less strongly than their S-timescale counterparts. In contrast, the structure of correct decision networks (e.g. H and C) was invariant across timescales. Note that S- and L-timescale interactions are nested, i.e., that the former are a subset of the latter. Given that we interpret L-timescale interactions to include interactions with more distant neurons, the invariance of the correct decision network structure between S- and L-timescales implies the existence of a network of local cortical interactions, rather than of interactions mediated by wider loops involving further away neurons, for performing the task correctly.

For a given S- or L-timescale, behavioral choice modulated network structure. For S-timescale networks, comparison of H- and M-trial networks showed the former had more links and larger subnetworks, suggesting that larger networks are beneficial for encoding correct detection of the target. The average link strength in C-trial networks was also greater than in F-trial network, suggesting that stronger links are beneficial for encoding correct rejection of the non-target. In contrast, for L-timescale networks, both the number of links and the sizes of subnetworks were smaller for correct choice than incorrect choice networks, while links remained stronger for correct choice networks. Together, these results indicate that aspects of network structure are key for encoding correct behavioral responses to target and non-target sounds during an auditory discrimination task. We next set out to examine this implication.

### Neuronal network structure encodes behavioral choice

Since the GC network structures for neurons with low *II*-peak latency strongly depended on behavioral choice, we sought to directly test if the network structures encode behavioral choice. Thus, we used the four GC network statistics as features for a support vector machine (SVM) trained to distinguish between correct (H or C) and incorrect (M or F) decisions. For comparison, we trained a similar classifier for networks of neurons that show high response rates, chosen regardless of the information content they carry. The GC network statistics of highly responsive neurons are reported in Figure 3 – Figure Supplement 1 and Supplementary Table S1. Of all low *II*-peak latency neurons, 30.21% were also identified as highly responsive neurons (see Methods for selection criterion). The neuronal network structure of S timescale networks for low *II*-peak latency neurons classified behavioral choice much more accurately than the network structure of highly responsive neurons (Fig. 3D, left bar plots). In contrast, the features of L timescale networks classified behavioral choice well for both low *II*-peak latency and highly responsive neuronal networks, though classifications for the former were more accurate (Fig. 3D, right bar plots). These results show that S timescale networks of low *II*-peak latency neurons better encode behavioral choice than those of highly responsive neurons and suggest that low *II*-peak latency neurons form a specialized group of neurons in A1.

One possibility is that strong choice predictivity from network interactions is not a special property of networks formed by neurons with *II* (i.e. neurons specifically carrying sensory information used for behavioral choice), but is also present in networks of neurons with either *SI* not used for choice or *CI* not related to the stimulus. That is, networks of exclusively stimulus-selective or choice-selective neurons, without intersection between the two signals, may also be strongly predictive of choice. To address this possibility, we compared the predictivity of low *II*-peak latency neurons to that of *SI* and *CI* neurons that did not have significant *II* (Fig. 4 and Table 2). Specifically, 20 neurons with either above-threshold peak *SI* or *CI* but sub-threshold peak *II* that had the shortest *SI*/*CI* peak latency were selected for GC network analysis in each session (N=8), also with at least 20 *SI*/*CI* neurons. Network structures of low *II*-peak latency neurons were more predictive than *SI*/*CI* neurons (Figure 4B). Furthermore, the network structures of neurons with the greatest *II* peak magnitudes (Fig. 3 Supp. 2 and Supp. Table S2) were also more predictive of behavioral choice than highly responsive neurons (Fig. 3 Supp. 2B). Our results thus suggest that the encoding of behavioral choice in the S-timescale network structure is specific to high *II* neurons, and it is not found as much in groups of neurons with either stimulus information not used for choice or choice information not related to the stimulus. Moreover, our results indicate that task-relevant information can be encoded not only by neuronal activity, but also in the structure of neuronal functional networks.

**Figure 4.**
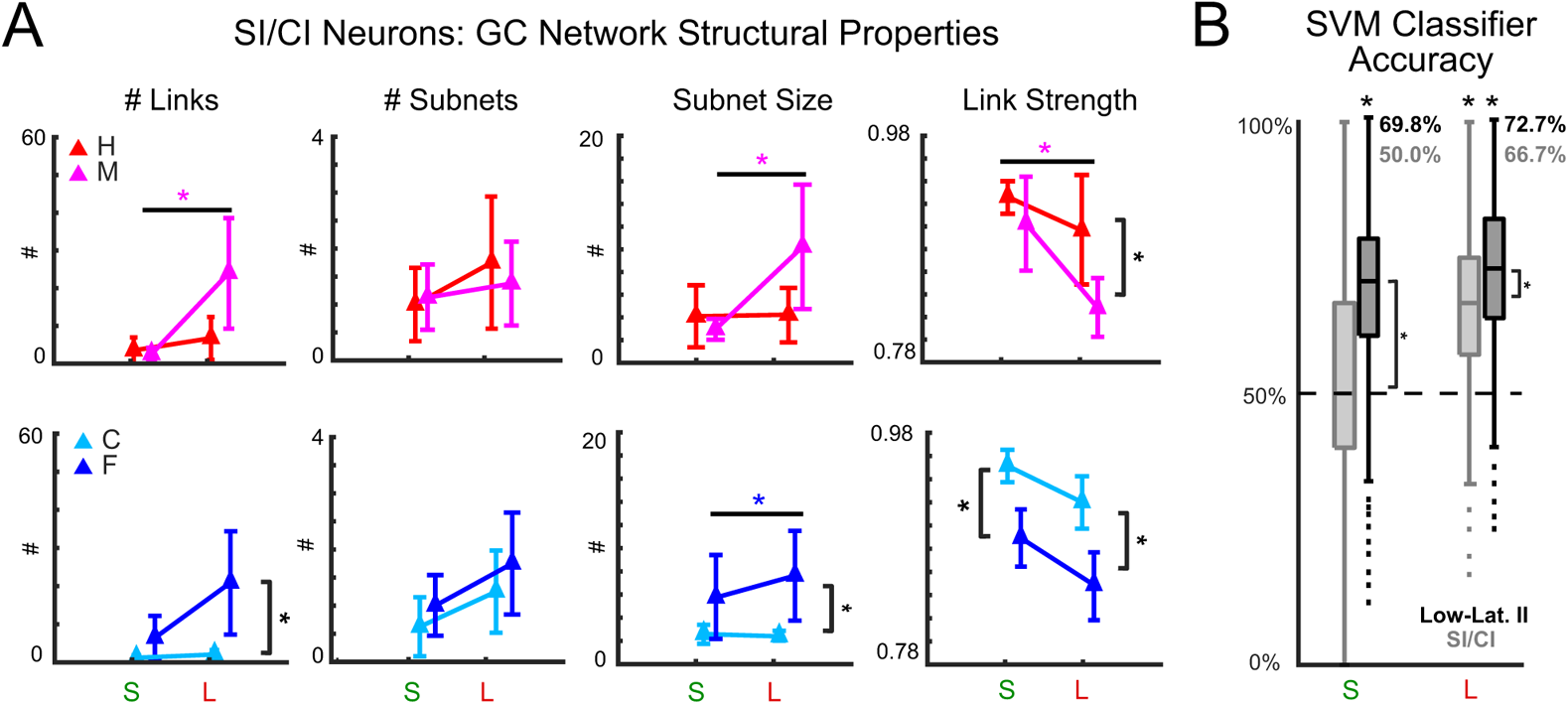
Network structure of neurons with either *SI* or *CI*, but not *II*, is less predictive than low-latency *II* neurons. **A.** Functional networks of short (S)- and long (L)-timescale interactions amongst neurons with high *SI* or *CI*, but not *II*, were estimated using Granger Causality (GC) analysis for each behavioral choice: Hit (H), Miss (M), Correct Rejection (C), and False Alarm (F). Four GC network statistics were analyzed: number of links, number of subnetworks, size of subnetworks, and statistical strength of links. Network statistics of *SI/CI* neurons are similar between behavioral choices at S-timescales, excepting link strength in C vs. F networks; additional differences in network statistics emerged at L-timescales. See also Table 2. **B.** Network statistics were used to train an SVM to classify into correct or incorrect decisions. Across timescale and selection of neurons—except *SI/CI* S-timescales—decisions were predicted significantly better than chance (p<0.001). S-timescale network structure of *SI/CI* neurons was decoded at chance-level accuracy, less than of low-latency *II* neurons (p<0.001), shown in Fig. 3. L-timescale network structure had higher decoding accuracy, but *SI/CI* neuronal networks were less accurately decoded (p<0.001). Thus, task-relevant stimulus information (*II*), rather merely than the combination of *SI* and *CI*, encodes behavioral choice in a tone discrimination task. Panel **A** shows means ± 2 SEM. Asterisks indicate statistically significant differences. Wilcoxon’s signed rank test (p<0.05) was used in **A**. A two-sample t-test (p<0.05) in **B** to compare neurons with low-latency *II*, and a one-sample t-test (p<0.05) to compare performance with chance decoding accuracy.

**Table 2.** Neurons with *SI/CI*, excluding *II* neurons: statistical comparisons of GC network structure across short (S) and long (L) timescales, and behavioral choice categories — hit (H), miss (M), correct-rejection (C), and false-alarm (F) – using Wilcoxon’s signed rank test (p<0.05). See also Figure 4A.

|  | # Links |  |  | # Subnets |  |  | Subnet Size |  |  | Link Strength |  |  |
| --- | --- | --- | --- | --- | --- | --- | --- | --- | --- | --- | --- | --- |
|  | S | L | p-value | S | L | p-value | S | L | p-value | S | L | p-value |
| H | 3.25 ± 1.810 | 6.50 ± 2.822 | 0.352 | 1.0 ± 0.327 | 1.75 ± 0.590 | 0.29 | 4.0 ± 1.334 | 4.14 ± 1.157 | 0.937 | 0.93 ± 0.007 | 0.90 ± 0.245 | 0.357 |
| M | 2.25 ± 0.861 | 23.88 ± 7.30 | **0.021** | 1.13 ± 0.295 | 1.38 ± 0.375 | 0.609 | 2.89 ± 0.455 | 10.0 ± 2.680 | **0.025** | 0.90 ± 0.021 | 0.83 ± 0.13 | **0.027** |
| p-value | 0.629 | 0.053 |  | 0.781 | 0.602 |  | 0.452 | 0.065 |  | 0.39 | 0.065 |  |
| C | 1.13 ± 0.516 | 2.0 ± 0.567 | 0.273 | 0.63 ± 0.263 | 1.25 ± 0.366 | 0.189 | 2.60 ± 0.40 | 2.40 ± 0.221 | 0.676 | 0.95 ± 0.007 | 0.92 ± 0.011 | 0.091 |
| F | 6.38 ± 2.885 | 20.88 ± 6.735 | 0.078 | 1.0 ± 0.267 | 1.75 ± 0.453 | 0.181 | 5.75 ± 1.840 | 7.64 ± 1.935 | 0.487 | 0.89 ± 0.012 | 0.85 ± 0.015 | 0.082 |
| p-value | 0.114 | **0.026** |  | 0.334 | 0.406 |  | 0.135 | **0.018** |  | **0.007** | **0.005** |  |

### The spatial extent of neuronal subnetworks is timescale-invariant during correct behavioral choices

Cortical neurons can interact over different spatial scales, thus functional networks might be composed of neurons in specific local regions or involve more dispersed and distant neurons. Since 2P imaging gives the exact spatial location of each neuron in a field of view, we sought to characterize how *II* neurons and functional networks thereof were distributed across the imaged area. We first studied if A1 L2/3 with *II* peaks in *peri* vs. *post-stimulus* time-windows were located in different regions in or if they were intermingled. We calculated the sum of the average distances of *peri-(Pe)* and *post-stimulus (Po)* neurons to their centroids (R_Pe_ and R_Po_, respectively) and compared the sum to the distance between the centroids (R_Pe - Po_). The distance between centroids was smaller than the spread of each set of neurons (Fig. 5A). Thus, *Pe* and *Po* neurons were heterogeneously distributed within each 2P imaging field of view. Since *Pe* and *Po* neurons were intermingled, this analysis suggests that localized information flow did not have intrinsic directionality from one subarea to another subarea during task performance.

**Figure 5.**
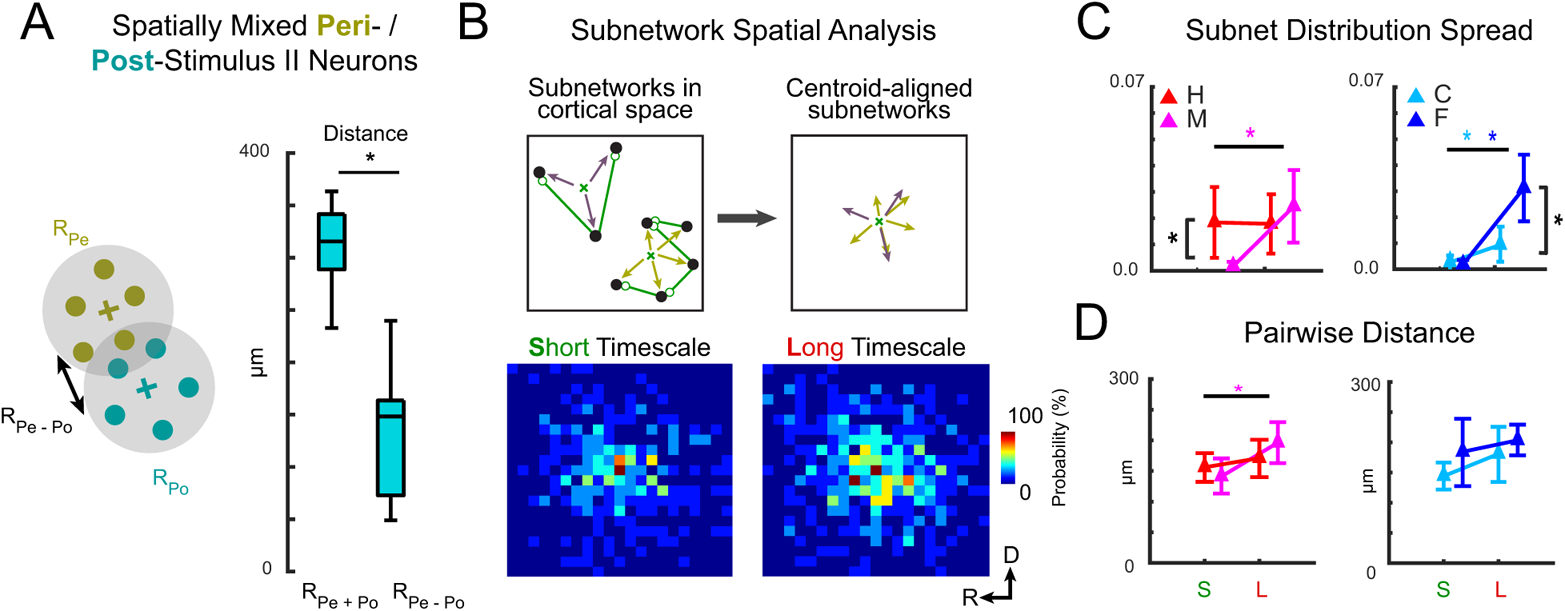
Neuronal subnetwork dispersion is timescale-invariant during correct behavioral choices. **A.** Neurons with peri-(*Pe*) and post-stimulus (*Po*) *II* peaks were spatially intermingled. The sum of average distances of *Pe* neurons to their centroid (R_Pe_) and of *Po* neurons to theirs (R_Po_) was smaller than the distance between centroids (R_Pe - Po_) (p<0.001), as determined by a two-sample t-test with threshold p<0.05. **B.** Subnetwork spatial distributions. Low *II*-peak latency neurons (black) that are linked (green) in groups isolated from others constitute subnetworks (top left). The locations of subnetworked neurons relative to their centroid were aggregated over all subnetworks (top right). The distributions of relative locations are shown as 2D histograms (25 µm x 25 µm bins) for S and L timescales (bottom left and bottom right). **C.** Determinant of spatial distribution covariance matrix. L-timescale C, M, and F trial subnetworks were more spatially dispersed than S-timescale subnetworks (M: p<0.001; F: p=0.002; C: p=0.014). For S timescales, H-trial subnetworks were more dispersed than M-trials (p=0.002), as were F-trial v. C-trial subnetworks for L timescales (p=0.003) **D.** Pairwise distances between linked neurons remained similar for S- vs. L-timescales, except for M-trials (p=0.047). Thus, subnetworks dispersed due to inclusion of more neurons. Panels **C** and **D** show mean ± 2 SEM. Asterisks indicate statistically significant differences. Comparisons were made using Wilcoxon’s signed rank test (p<0.05).

Though neurons carrying task-relevant information were not located in spatially segregated positions, their functional networks could differ in spatial organization by timescale or behavioral choice. Hence, we analyzed how neuronal subnetworks were dispersed across the imaged area by computing the vector distances of subnetworked neurons to the subnetwork centroid (Fig. 5B, top schematic; see also Methods). Subnetworks of L-timescale interactions tended to be more spatially dispersed than S timescale subnetworks (Fig. 5B, bottom subpanels), and this effect was confirmed by computing the determinant of the distance vector covariance matrix (Fig. 5C). The dispersion of M- and F-trial subnetworks were larger for L- than S-timescales. Additionally, L-timescale subnetworks were more dispersed than S-timescale subnetworks for C-trials, though to a lesser degree than in M- and F-trials. Differences in dispersion were also observed between H and M trials for S-timescale subnetworks and between C and F trials for L-timescale subnetworks. In summary, we find that GC neuronal networks became the most spatially dispersed across A1 for L-timescale interactions and during incorrect behavioral choices. These results suggest that correct choices are associated with subnetworks whose spatial spread do not vary by timescale.

We next asked if differences in the dispersion of subnetworks across timescales were due to greater distances between linked neurons, rather than the inclusion of additional neurons. Thus, we analyzed network spatial dispersion using the average pairwise distance between linked neurons, i.e., the average link length (Fig. 5D). Together we found that except for M-trials, GC link lengths were stable across timescales, indicating that the greater subnetwork spatial dispersion for L-timescale interactions was more likely due to the inclusion of additional neurons than an increased distance between linked neurons. These results suggest that correct choices are associated with spatially stable compact subnetworks while incorrect choices involve activity spread to additional neurons.

### Networked neurons communicate redundant information

Our results show that auditory task-relevant information is transiently encoded by individual neurons in A1 L2/3 yet sustained over longer periods by sequential encoding between functionally linked neurons. A functional link between neurons suggests that task-relevant information is transmitted from one neuron to another. This would create a population code whose information content is reverberated redundantly across neurons because the same information is shared by different neurons as a result of the transmission. Redundancy would reduce the amount of total information that the network can accumulate over time, but would also generate a more robust information representation that is distributed across neurons, more persistent over time, and could facilitate task-relevant information readout for behavioral choice (Runyan, Piasini et al. 2017).

To investigate the nature of information present in the functional networks, we measured information redundancy (Pola, Thiele et al. 2003, Schneidman, Bialek et al. 2003) between GC-linked neurons. We used a normalized redundancy index defined as the information carried jointly by two neurons minus the sum of the information that each carried independently, normalized with respect to the total information carried by the two neurons jointly. Neurons share redundant information when the redundancy index is negative; that is, together they carry less information than the sum of the information they carry separately. The value of the normalized redundancy index indicates the fraction of total joint information that is shared among the two neurons.

For *SI, CI,* and *II*, we computed redundancy using the information that each neuron carried at its peak time of *II*, for each pair of the 20 different neurons selected in every session for GC analysis. Normalized redundancy between pairs of neurons with a S-timescale GC link was compared to those with no GC link (Fig. 6A, left panel). For S-timescales, we found that information shared between pairs of neurons was redundant (i.e., neurons together carried less information than the sum of their individual information), implying that neurons within A1 shared part of the information they transiently carried at different times. Normalized redundancy was much larger for *II* than *CI* and *SI* (Fig. 6A, left bar plots vs. middle and right bar plots). This means that neurons shared more of the behaviorally-relevant, than the behaviorally-irrelevant, portion of the *SI* that they carried. Importantly, the difference between normalized redundancy for GC-linked vs GC-unlinked neurons was much larger for *II* than for *SI or CI* (Fig. 6A, red vs. black bar plots), reinforcing the interpretation that S-timescale GC links mediate the exchange of behaviorally relevant sensory information.

**Figure 6.**
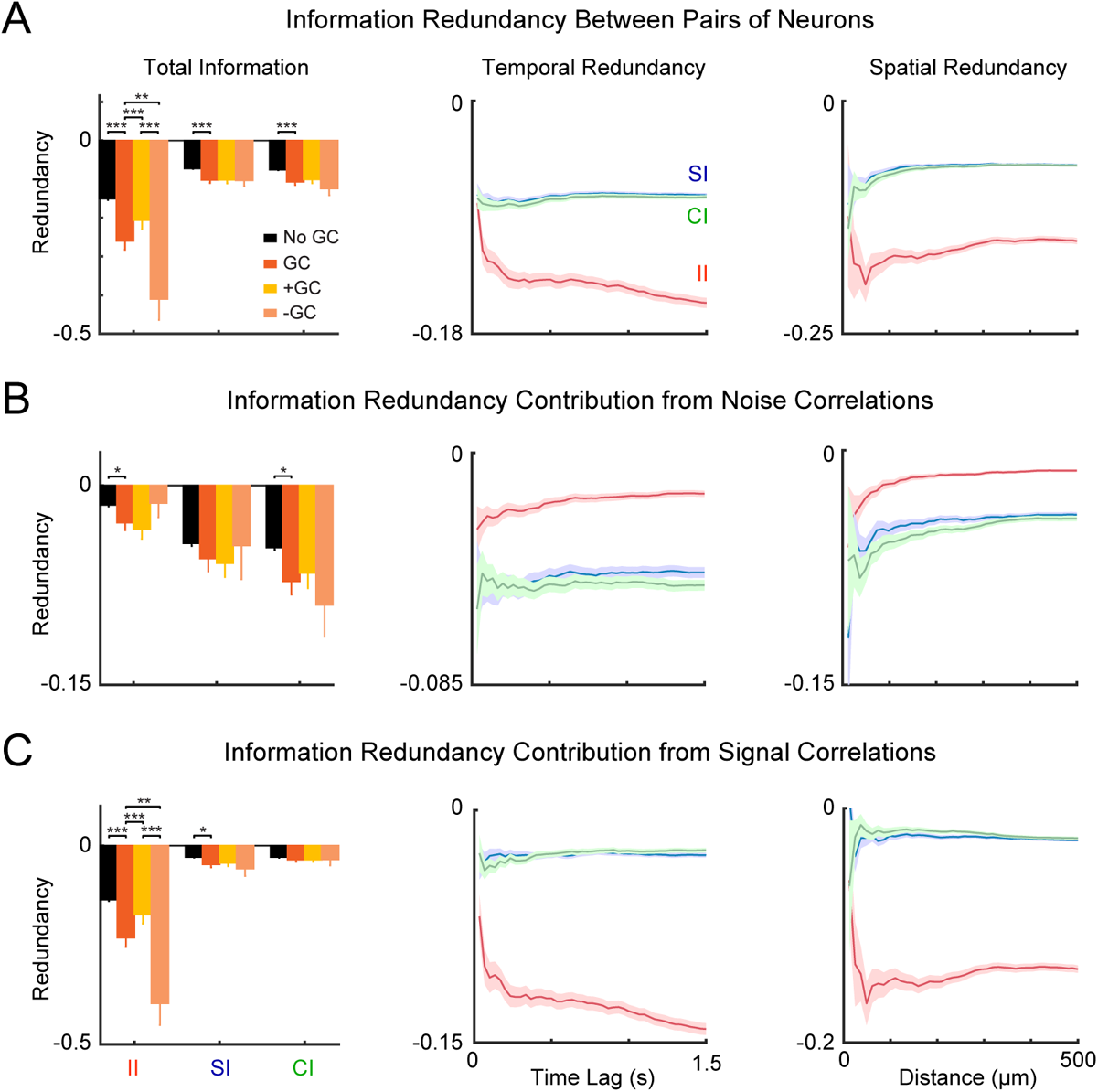
GC-linked neurons share redundant information. **A.** The left panel shows normalized time-lagged redundancy computed for GC-linked neurons (red), that were further split into positive (yellow) or negative (pink), and GC-unlinked pairs of neurons (black). GC-linked pairs of neurons carried more redundant information than GC-unlinked pairs of neurons (*II, SI, CI*). Pairs of neurons connected with negative GC-links carried more redundant information related to *II*. Normalized redundancy across time-lagged neuronal activity (middle panel), and as a function of the Euclidean distance (right panel) between pairs of both GC-linked and GC-unlinked neurons. **B, C**. We breakdown information redundancy contributions from noise (**B**) and signal (**C**) correlations in the different groups of GC-linked and GC-unlinked neurons. In case of noise correlations, we report higher values of redundant information in GC-linked pairs of neurons (*II* and *CI*), while for signal correlations there was a significant difference for *II* and *SI*. Pairs of neurons connected with negative GC links showed higher redundancy index than pairs of neurons linked by positive GC links for signal correlations (*II*). Solid lines and shaded regions represent mean±SEM (N=#pairwise links between neurons of a given group), respectively. Statistical comparisons between groups are made with a two-sample t-test (*p<0.05; **p<0.01; ***p<0.001).

GC links can be positive or negative valued, reflecting functionally facilitative or suppressive interactions, respectively (Francis, Winkowski et al. 2018, Sheikhattar, Miran et al. 2018). We found that negative GC links had a much larger effect on redundancy, suggesting they mediate more *II* exchange than positive links (Fig. 6A, orange vs. yellow bar plots). These results might indicate a mediating role of inhibitory circuits in task-related network activity (Kuchibhotla, Gill et al. 2017).

Sorting the normalized redundancy with respect to the *II* peak time lags (Fig. 6A, middle panel) revealed that *II* redundancy varies across time lags with an overall increasing trend (from -0.1 to -0.15). This indicates that redundant information persists over the trial.

Previous studies showed that nearby cells typically interact redundantly (Nirenberg, Carcieri et al. 2001, Reich, Mechler et al. 2001, Chechik, Anderson et al. 2006). We thus investigated how redundant information spreads spatially for *II, SI*, and, *CI* in our experiments by plotting the time-lagged redundancy as a function of the Euclidean distance between pairs of neurons (Fig. 6A, right panel). We found a peak of redundant interaction for the total *II* at a distance of ~50 *μm* (*II* = −0.1971 ± 0.0187) which then reached a plateau at ~320 *μm* (*II* = −0.1499 ± 0.0039), followed by a distance-independent trend. *SI* and *CI* were similarly redundant and reached a plateau at ~208 *μm* (*SI* = −0.0691 ± 0.0019, *CI* = −0.0709 ± 0.0019. Together, these results suggest that GC links indicate redundant communication of behaviorally-relevant stimulus information, and that redundant neurons are located in close proximity of each other.

If information is passed from one neuron to another, we would expect more information redundancy between two GC-linked neurons due to similarity in their shared “signal” (the covariations of activity that reflect similarity in tuning to stimulus, choice or intersection signals) than due to correlation in their shared “noise” (the covariations of activity that are unrelated to the tuning to stimulus, choice or intersection signals). We separated and quantified the contribution of signal similarity and noise correlation to the total redundancy shared by two neurons using the information breakdown (Pola, Thiele et al. 2003, Schneidman, Bialek et al. 2003). Consistent with the hypothesis that a GC link transmits mainly behaviorally-relevant stimulus information, we found that GC-linked neurons in S-timescale networks had much greater *II* redundancy due to signal correlations than due to noise correlations (Figs 6B,C). The prevalent contribution of signal correlation to redundancy was specific to *II* and not found as much in SI and CI. Importantly, GC-linked neurons had much greater contributions of signal correlations to *II* redundancy than GC-unlinked neurons (Fig 6C), further corroborating that GC links mediate exchange of *II* across linked neurons. Finally, we found that the contribution to *II* redundancy of noise correlations, though small, was higher for GC-linked neurons than for GC-unlinked neurons (Fig 6B). This result is consistent with the notion that connected neurons may additionally share information unrelated to the task variables. Notably, the contribution of noise correlation to *CI* redundancy was largest (that is, the most negative) (Fig. 6B, bar plots in the *CI* column), suggesting that pre-existing intrinsic network dynamics carrying task-unrelated signals also contribute some to *CI*.

The contribution of signal correlations to *II* redundancy peaked at short time lags and distances (Fig. 6C, middle and right panels), whereas the contribution of signal correlation to both *CI* and *SI* redundancy was small and did not vary over time and space (Fig. 6C, middle and right panels). The contribution of noise correlations was relatively stable over time and space for all information types (Fig. 6B, middle and right panels). This suggests a partial decoupling of the spatial and temporal scales information transmission for task-specific vs task-unspecific signals, with sensory information relevant for choice being shared more at shorter time scales than other kinds of information.

We finally examined the contributions of signal and noise correlations to *II* redundancy on L-timescale networks (Fig. 6 – Figure Supplement 1). While the contribution of signal and noise correlation to *SI* and *CI* redundancy showed similar modulation trends, they exhibited proportionally smaller variations between GC-linked and GC-unlinked neurons, and the contribution of signal correlations to redundancy was less strong than for S timescales.

Together, these results further corroborate the view that short timescale GC links are those principally mediating the communication of behaviorally-relevant stimulus information between neurons.

## DISCUSSION

In this study we found that during the performance of an auditory discrimination task, individual neurons in A1 L2/3 transiently and differentially carried information about the stimulus (*SI*), behavioral choice (*CI*), or both (*II*) for hundreds of milliseconds and that task-relevant information was sustained across the duration of a three-second trial by sequential propagation of *SI, CI,* and *II* in functionally connected neuronal populations. Furthermore, we identified a subpopulation of low *II*-peak latency neurons, which formed functionally connected networks whose structure could predict behavioral choice better than a subpopulation of highly responsive neurons. Our findings suggest that the spatiotemporal structure of functional connectivity between low *II*-peak latency neurons in A1 L2/3 may form a neural population base for sustained representation of task-relevant information.

A1 L2/3 contains a diverse population of neurons with differing functional connectivity (Meng, Winkowski et al. 2017) We here find that the bandwidth of high II neurons is lower than that of other neurons. This suggests that these neurons might be part of a class of A1 L2/3 neurons which receive L4 inputs, have limited integration across the tonotopic axis and lower bandwidth (Meng, Winkowski et al. 2017)

### Task Relevance of Short vs Long Timescale Neuronal Interactions

The nested parameterization of L- and S-timescale interactions allowed us to differentiate between solely S-timescale influences vs. additional L-timescale influences in functional networks. Comparing L- vs. S-timescale networks showed that correct choice L-timescale networks consisted of fewer but stronger links that were mostly S-timescale influences. Thus during correct choices, GC active networks are constrained. In contrast, incorrect choice networks are characterized by a mixture of both S- and L-timescale links, and by an increased network size due to recruitment of additional spatially distant neurons. These additional L-timescale links likely reflect the local effects of slower interactions with distant neurons, perhaps reflecting non-sensory task-related interaction, such as error-signaling or deviance detection (Chen, Helmchen et al. 2015, Khouri and Nelken 2015, Steinmetz, Zatka-Haas et al. 2019, Stringer, Pachitariu et al. 2019, Parras, Casado-Roman et al. 2021). Since subnetworks during correct trials were timescale-invariant, this suggests that the influence of more distant neurons is suppressed when correct decisions are made, leaving predominantly S-timescale interactions. Such suppression could be mediated by inputs to A1 which can activate inhibitory circuits (Fritz, David et al. 2010, Winkowski, Bandyopadhyay et al. 2013, Winkowski, Nagode et al. 2018, Liu, Xin et al. 2021).

### Population Coding via Reverberation of Redundant Information in Networks

We found that there was high redundancy between the behaviorally-relevant stimulus information carried at the time of information peaks between pairs of low *II*-peak latency neurons. This spatio-temporal redundancy was higher between pairs of neurons that were functionally GC-linked based on S-timescale interactions, suggesting that the GC link may reflect the transfer of behaviorally-relevant information from one neuron to another. Redundancy has been traditionally viewed as a negative feature of population coding that should be reduced or eliminated, based on theories of efficient coding (Attneave 1954, Barlow 1961, Nigam, Pojoga et al. 2019), and on the often implicit assumption that information is read out optimally, so that higher neural information corresponds to higher discrimination performance (Gold and Shadlen 2001). However, other studies have proposed that high values of spatiotemporal redundancy might benefit task performance by facilitating biophysical signal propagation (Alonso, Usrey et al. 1996, Salinas and Sejnowski 2001, Valente, Pica et al. 2021), thereby making the task-related signals available for longer and storing sustained activity more robustly (Runyan, Piasini et al. 2017), and facilitating the behavioral readout of the sensory signal.

These previous studies highlighting the role of redundancy in behavioral readout, however, concentrated only on the average strength of pairwise noise correlations and therefore could not shed light on the network structure underlying correct decisions. Here, we provided significant new findings about the network-level structure of behaviorally relevant information sharing and of correct perceptual decisions. We found that higher redundancy in GC-linked neurons was accompanied by (1) a higher number of links, (2) larger subnetworks in correct target detection, and (3) stronger links in correct target rejection. Thus, together, redundancy and functional connectivity analyses suggest that performing well on an auditory discrimination task may require a certain amount of information to be at least temporarily reverberated in the spatiotemporal structure of neuronal networks during task performance. This might explain why we found that the redundancy was larger for behaviorally-relevant (intersection) than behaviorally-irrelevant sensory information. In other words, information that does not reverberate in an appropriately structured neuronal network may not be useful for behavior.

In summary, here we employed an information theoretic framework to analyze neural activity during a pure-tone frequency discrimination task and quantified the encoding of task-relevant information by functionally connected neurons. Our results show that task-relevant information is transmitted sequentially across individual neurons in primary auditory cortex and is sustained for longer periods of time within compact neuronal networks.

## METHODS AND MATERIALS

### CONTACT FOR REAGENT AND RESOURCE SHARING

Further information and requests for resources should be directed to and will be fulfilled by the Lead Contact, Patrick O. Kanold.

### EXPERIMENTAL MODEL AND SUBJECT DETAILS

All procedures were approved by the University of Maryland Institutional Animal Care and Use Committee. We used N=9 mice (3 female, 6 male) F1 offspring of CBA/CaJ strain (The Jackson Laboratory; stock #000654) crossed with transgenic C57BL/6J-Tg(Thy1-GCaMP6s)GP4.3Dkim/J mice (Dana, Chen et al. 2014)(The Jackson Laboratory; stock #024275) (CBAxThy1), 8–24 weeks old, in 35 total experiments. We used the F1 generation of the crossed mice because they have good hearing into adulthood (Frisina, Singh et al. 2011). Each mouse was tested once per day over multiple days. The mice were trained to perform the task before collecting 2P data during task performance. Mice were housed under a reversed 12 h-light/12 h-dark light cycle and trained during the dark cycle.

## METHOD DETAILS

### Auditory task

We designed a pure-tone frequency discrimination task that used behavioral response-timing rules to induce well controlled behavioral responses in mice. Each mouse was first trained on a positive reinforcement tone detection task, with water used as a rewarding stimulus, as done previously (Francis, Winkowski et al. 2018). We then trained the mice on the frequency discrimination task. Each trial began with 1 second of silence, followed by a 55 dB SPL amplitude modulated (8 Hz) tone presented for 1 s. The target tone frequencies were 7 and 9.9 kHz. The non-target frequencies were 14 and 19.8 kHz. The tone frequency was randomized across trials. The tone was followed by 2 s of silence, and a random 5-9 s inter-trial interval (ITI). The tone was presented during every trial of task-performance, and the mice were trained to lick a waterspout 0.5 s after the onset of a target tone to receive a water droplet reward, and to avoid licking the waterspout after a non-target tone. Each trial’s behavioral response was categorized as a hit (licking after target onset), miss (no licking after a target), false alarm (licking after non-target onset), or correct rejection (no licking in response to a non-target). Incorrect behavioral responses were punished with an 8 s time-out added to the ITI. Mouse health was monitored daily by a skin turgor test and checking that body weight remained above 80% of the initial off-study weight.

### Imaging

Chronic window implantation, widefield imaging, and 2-photon imaging, were performed as previously (Francis, Winkowski et al. 2018). In brief, a chronic imaging window was implanted over a 3 mm craniotomy over auditory cortex. For widefield (WF) imaging, neuronal activity was quantified by comparing fluorescence during the stimulus versus the silent pre-stimulus baseline, resulting in a response amplitude (ΔF/F). After visualizing wide-field tonotopic maps, a site was selected for 2-photon (2P) imaging in primary auditory cortex (A1) for each mouse. For each 2P imaging site, we determined the frequency selectivity (best frequency [BF]) of individual neurons during passive trials, i.e., trials when the mouse sat quiescently hearing tones without doing an auditory task. BFs were determined from neuronal responses to 55 dB SPL pure tones ranging from 4 - 56.6 kHz. We used a scanning microscope (Bergamo II series, B248, Thorlabs) coupled to a pulsed femtosecond Ti:Sapphire 2-photon laser with dispersion compensation (Vision S, Coherent). The microscope was controlled by ThorImageLS software. The laser was tuned to λ = 940 nm. The field of view was 370 x 370 μm. Imaging frames of 512×512 pixels (pixel size 0.72 μm) were acquired at 30 Hz by bidirectional scanning of an 8 KHz resonant scanner.

A different set of neurons was imaged for each experiment. Using an average field of view from each experiment, the somatic centers of putative neurons were manually localized and stored. A ring-like region of interest (ROI) was cropped around the cell center using the method described in Chen, Wardill et al. (2013). Overlapping ROI pixels (due to closely juxtaposed neurons) were excluded from analysis. For each labeled neuron, a raw fluorescence signal over time was extracted from somatic ROIs. Pixels within the ROI were averaged to create individual neuron fluorescence traces, F_C_(t), for each trial of the experiment. Neuropil fluorescence was estimated for each cellular ROI using an additional ring-shaped ROI, which began 3 pixels from the somatic ROI. Pixels from the new ROI were averaged to obtain neuropil fluorescence traces, F_N_(t), for the same time-period as the individual neuron fluorescence traces. Pixels from regions with overlapping neuropil and cellular ROIs were removed from neuropil ROIs. Neuropil-corrected cellular fluorescence was calculated as 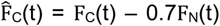. Only cells with positive values obtained from averaging 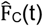 across time were kept for analysis, since negative values may indicate the dominance of neuropil contamination. ΔF/F was calculated from 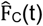, for each neuron, by finding the average F taken from the silent baseline period, subtracting that value from subsequent time-points, then dividing all time-points by the baseline F. All images were processed using Matlab (The Mathworks) using our prior methods (Francis, Winkowski et al. 2018).

### Computation of stimulus, choice and intersection information for single neurons

We quantified stimulus encoding and choice decoding using Shannon’s mutual information. We computed mutual information carried by neurons either about stimulus category S (low vs high frequency tones), and about the behavioral choices C (lick vs. no-lick). Mathematically, the mutual information for two discrete random variables is defined as follows (Cover and Thomas 1991, Quian Quiroga and Panzeri 2009):

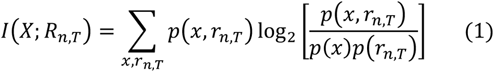

where *X* = *S, C* denotes the set of task variables, either stimuli S or choices C. *R_n,T_*) is the set of responses of the n^th^ neuron measured at a given time-point *τ*. *p*(*x*, *r_n,τ_*)*p*(r_n,T_) denotes the joint probability of observing in a given trial a value × for the stimulus or choice variable and a value *r_n,T_* for the activity of the n^th^ neuron at a given time-point τ. *p*(*x*) = ∑_*r*_*p*(*x*, *r_n,T_*), and *p*(*r_n,T_*) = ∑_*x*_*p*(*x*, *r_n,T_*), are the marginal probabilities.

Details of the calculation were as follows. To compute the neural responses *r_n,T_*, we first inferred the relative spiking activity from the fluorescence traces with a first-order autoregressive model using the CaImAn algorithm (Vogelstein, Packer et al. 2010, Giovannucci, Friedrich et al. 2019). We obtained similar results with a second-order auto-regressive model. We then adopted a sliding window approach to estimate the spike rates at different times during the perceptual task. We thus averaged the spiking activities over the imaging frames enclosed within a sliding window of ~300ms, which shifts in time-steps of ~30ms from 1 s before the stimulus onset until the end of the trial (3 s after the stimulus onset). Finally, for any time-point, neural responses were mapped to 0 if no spike occurred at all within the sliding windows of ~300ms, and 1 otherwise. At each time-point, we then computed mutual information from Eq (1) above with the Information Breakdown Toolbox (Magri, Whittingstall et al. 2009). To correct for the bias due to the finite number of trials one can estimate the shuffled information and subtract it from the time-course information. We found that even after correcting for this bias there were still non-zero values of both stimulus and choice information in the pre-stimulus period. They arose only as an artifact of spike-inference since bias-corrected pre-stimulus information was absent when computed from Δ*F*/*F* traces. Thus, we accounted for both the bias and the artifact by subtracting the overall information with respect to the baseline information computed in the pre-stimulus interval. Moreover, and importantly, with both the deconvolved and ΔF/*F* signals the pre-stimulus choice information was always very small and equal to the pre-stimulus stimulus information, meaning that there was no additional stimulus-unrelated choice information (e.g. some internal bias signal) in pre-stimulus neuronal activity. Thus, we estimated the average information in the pre-stimulus interval and subtracted it from the total amount of information computed in the interval after the tone-onset.

We then computed a further information quantity, called intersection information *II*(*S*, *R*, *C*), defined as the part of sensory information used for behavioral read out at the single-trial level (Pica, Piasini et al. 2017). Mathematically, we defined it as follows (Panzeri, Harvey et al. 2017, Pica, Piasini et al. 2017):

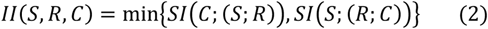

Where the term *SI*(*X*: {*Y; Z*}) denotes the *shared information* about the target X, that is carried from any of the sources Y and Z in a redundant way (Williams and Beer 2010, Pica, Piasini et al. 2017). *II*(*S*, *R*, *C*), has the important property of being bounded by stimulus *I*(*S*; *R*), and choice information *I*(*C*; *R*), and the mutual information between stimulus and choice *I*(*S*; *C*) (Pica, Piasini et al. 2017). The optimization problem in Eq.(2) was solved numerically with the BROJA algorithm (Makkeh, Theis et al. 2018). We used the same bias correction as did for stimulus and choice information by quantifying the average intersection information evaluated in the pre-stimulus interval and subtracting it from those values of intersection evaluated in the later interval after the tone onset.

To select individual neurons with significant information, we used a non-parametric permutation test (creating a null hypothesis distribution of information values obtained randomly shuffling across trials the stimulus-response or choice-response associations), and we set a mild threshold of p < 0.1 rather than a stronger threshold at p < 0.05 because the former enabled us to select 12 (rather than 7) experimental sessions with at least 20 *II* neurons for Granger causality network analysis. This criterion gave us a much higher statistical power for network analysis at the relatively small price of risking adding a small percentage of neurons (up to 5% of the total selected) with possible weakly significant or spurious information to inferred networks. Note that we constructed the null hypothesis distribution selecting for each random permutation the maximum information over all time windows of the permuted information values. Therefore, the so obtained p values are already corrected for multiple comparisons across time bins.

### Granger Causality Analysis

Granger causality (GC) analysis evaluates the predictive influence of the past activity of one neural process on present activity of another. GC analysis was performed similarly as in our previous work (Francis, Winkowski et al. 2018) by fitting sparse vector autoregressive (VAR) models to the ensemble neural responses (ΔF/F), calculating an unbiased GC measure for each potential link, and characterizing the GC link strengths using Youden’s J-statistics following false discovery rate control at a rate of 0.001. We highlight here three key differences from previous analysis regarding model estimation, modelling history-dependency, and neuron selection, and refer the reader to (Francis, Winkowski et al. 2018) for a recapitulation of the remaining details.

In order to estimate GC network connectivity amongst larger networks, the maximum likelihood problem in (Francis, Winkowski et al. 2018) is solved, employing the Orthogonal Matching Pursuit (OMP) algorithm (Cai and Wang 2011, Zhang 2011) to fit sparse VAR models rather than ℓ_1_-regularisation. OMP enables the sparsity of the estimated parameter vector—i.e. the number of non-zero parameters—to be controlled, thus mitigating model overfitting more robustly. The sparsity level of each VAR model is obtained by cross-validation. The set of non-zero parameters, called the model support set, is iteratively selected: at each iteration, a new parameter with the greatest contribution to the residual estimation error is added to the support and maximum likelihood estimation is performed over the updated support set.

The neural responses of a set of *C* neurons, indexed by *c* = 1, …, *C*, are denoted by 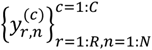, where *n* = 1, …, *N* and *r* = 1, …, *R* index time bins and trial repetitions, respectively. The covariates of the VAR model of each neural response incorporate the self- and cross-histories of activity over an integration window of *L* samples within which neuronal interactions are assumed to occur. The integration window is subdivided into *M* non-overlapping windows of lengths *{W_m_}_m=1:M_*. The average activity of neuron (*c*) in the *m*-th window lag with respect to time bin *n* and trial *r* is given by

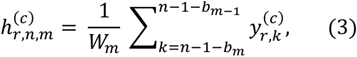

where 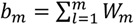 and *b*_0_ = 0. The collection of history covariates 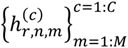 comprises the 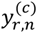 regressors of 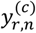. Interactions between neurons over short timescales (S) are modelled with an integration window of *L* = 7 lags with *M* = 3 subdivisions with window lengths *{W_m_}_m=1:M_* = {2^*m*−1^|_*m*=1:*M*_. Long timescale (L) interactions are modelled by instead using a cross-history integration window of length *L* = 31 lags with *M* = 5 subdivisions whose window lengths are similarly defined. S and L timescale interactions thus respectively correspond to 233 ms and 1033 ms windows of effective history. It is clear from the parameterization that the S and L interactions are modelled in a nested fashion. We validate this approach by simulating a 10 neuron network consisting of both S- and L-timescale links (see Figure 3 – Figure Supplement 3). Employing the L integration window for GC analysis, we are able to correctly identify all L- and S-timescale interactions; however, using the S integration window, while the S-timescale links are correctly identified, the L-timescale links are expectedly discarded, thus corroborating the sensitivity and specificity of our proposed inference framework.

Twenty neurons were analyzed from each 2-photon experiment. Analyzing a subset of fixed size avoids intersession variations in the number of recorded neurons that could affect analyses. The total number of model parameters, *M* · *C*, needs to be much smaller than the total number of samples, *R* · *N*, for reliable model estimation. We use at most *M* = 5 subintervals and per trial used the *N* = 105 time samples of the response after stimulus onset; we calculated *C* = 20 to be the maximum number of neurons that satisfies this condition, conservatively assuming at minimum *R* = 10 trials per session of each behavioral choice category. In our main results, 20 *II* neurons with the lowest *II*-peak latency in each experiment (N=12) in which at least as many *II* neurons were identified. Highly active neurons in each 2-photon experiment (N=34) were selected per behavioral choice category. The neural response of the *c*^th^ neuron at the *n*^th^ time index of the *r*^th^ repeated trial of a behavioral category, 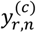, is normalized 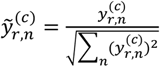. The 20 neurons with the smallest trial-averaged variances of the normalized responses, 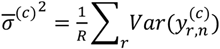, were selected.

### Decoding behavioral choice from GC network structure

To test if network structures encode behavioral choice, we trained classifiers on four GC network statistics — number of links, number of subnetworks, size of subnetworks, and statistical strength of links — to distinguish correct (Hit or Correct Rejection) and incorrect (Miss or False Alarm) decisions. Feature vectors consisting of these statistics were compiled for each behavioral choice network estimated from each experiment. We trained a linear support vector machine (SVM) on a randomly selected 75% of the feature vectors, with the remaining 25% used to evaluate prediction accuracy. This procedure was repeated 2000 times, each with a new randomized partition of feature vectors, to characterize the distribution of average classification accuracy.

### Spatial Distribution of GC subnetworks

To investigate the spatial scales over which functionally linked neurons interact, we leveraged the spatial location of individual neurons available in 2P imaging to analyze how subnetworks were distributed across the imaged cortical area. To this end, the locations of subnetworked neurons relative to their centroid were obtained as follows. For a subnetwork of *R* neurons with positions 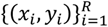, we compute their locations relative to the subnetwork centroid, 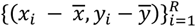, where 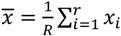 and 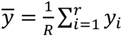 The relative locations are compiled over all subnetworks to yield an empirical distribution. The covariance matrix of the distribution describes the spatial spread of subnetworks; its determinant is used as a comparative statistic to quantify differences is the spatial dispersion of subnetworks across conditions.

### Computation of the spatio-temporal information redundancy and of its breakdown in the contribution of signal and noise correlations

To quantify the spatio-temporal redundancy of information we used a normalized redundancy index defined as the information carried jointly by two neurons minus the sum of the information that each carried independently, normalized with respect to the total information carried by the two neurons jointly (Pola, Thiele et al. 2003, Schneidman, Bialek et al. 2003):

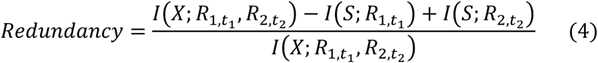

For each of the two neurons, we selected activity at the time *t_1_,t_2_* of their peak information. Thus, there was often a lag between the activities of the neurons considered for redundancy and it should thus be considered a measure of spatio-temporal redundancy. The information carried by each neuron individually was computed as defined in section “Definition of stimulus, choice and intersection information for single neurons”. The joint time-lagged stimulus and choice information was computed from the Shannon formula as follows:

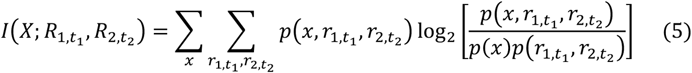

Notations are as in Eq (1) , with now *p*(*X*, *r*_1,t_1__,*r*_2,t_2__)) denoting the probability of observing in a given trial a value × of a the behavioral variable (stimulus category or choice) and a joint response *p*(*X*, *r*_1,t_1__,*r*_2,t_2__ of the two neurons at times *t_1_,t_2_* respectively. Intersection information was computed from Eq (2) (Pica, Piasini et al. 2017), but using the joint response *p*(*X*, *r*_1,t_1__,*r*_2,t_2__ as the neural response variable. We broke down the joint information into components associated to noise and signal correlations (Pola, Thiele et al. 2003, Schneidman, Bialek et al. 2003, Magri, Whittingstall et al. 2009). We computed a “shuffled” joint distribution by combining at random responses of the two neurons from different trials of the same behavioral (same stimulus category for stimulus information; same choice for choice information; same stimulus category and choice for intersection information). This shuffled joint distribution has the same value of marginal probabilities of both neurons but no within trial “noise” correlations. We estimated the joint information in 100 shuffled joint distributions and considered the mean value as the shuffled joint information. The contribution of signal correlation to redundancy was the difference between the joint information carried by the shuffled joint probability and the sum of the two individual information values (Pola, Thiele et al. 2003, Schneidman, Bialek et al. 2003, Magri, Whittingstall et al. 2009). The contribution of noise correlation to redundancy was the difference between the true value of joint information and the joint information carried by the shuffled joint probability (Pola, Thiele et al. 2003, Schneidman, Bialek et al. 2003, Magri, Whittingstall et al. 2009). The sum of the signal and noise component of redundancy equals redundancy. Like for the full redundancy, the value of its signal and noise components was normalized by dividing by the value of the true joint information.

### Quantification and statistical analysis

Unless noted otherwise, statistical comparisons were performed using a bootstrap t-test with 10000 iterations or a Kolmogorov–Smirnov test (KS-test), for both one- and paired-sample tests. Kruskal-Wallis tests were used when there were >2 groups being compared. We used a Bonferroni correction for multiple comparisons. All mean values are reported with 2 standard errors of the mean, unless noted differently.

### Data and software availability

The dataset and software are available upon request.

## ACKNOWLEDGEMENTS

The authors would like to thank Menatallah Mohamed for surgical assistance. Supported by NIH RO1DC9607 (POK), U01NS90569 (POK), P01AG55365 (POK), U19NS107474 (POK, BB, SP), R21DC017829 (NAF) and NSF ECCS1807216 (BB).

## SUPPLEMENTARY FIGURES

**Figure 2 – Figure Supplement 1.**
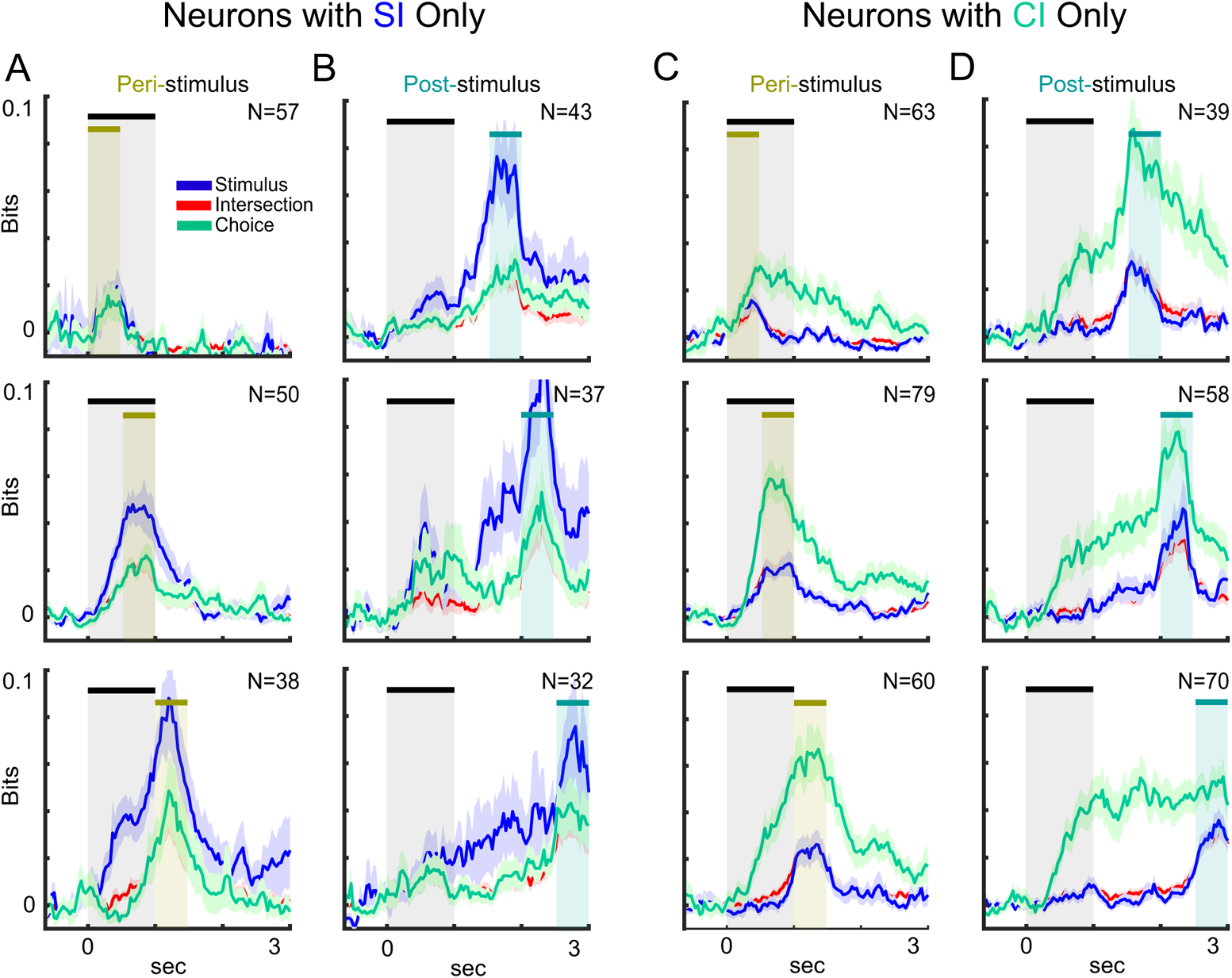
Time-course of *SI*, *CI*, and *II* averaged over neurons that carried stimulus information (*SI*) only in **A**-**B** and choice information (*CI*) only in **C**-**D**. As in Figure 2 in the manuscript, we quantified the *SI*, *CI*, and *II* in six separate stages of the behavioral task, which account for the peri-stimulus (0-1.5 s) and the post-stimulus intervals (1.5-3 s) shown by the shaded regions. Error bars show one standard error of the mean (SEM; N=#neurons with SI, CI peaks within the stage).

**Figure 3 – Figure Supplement 1.**
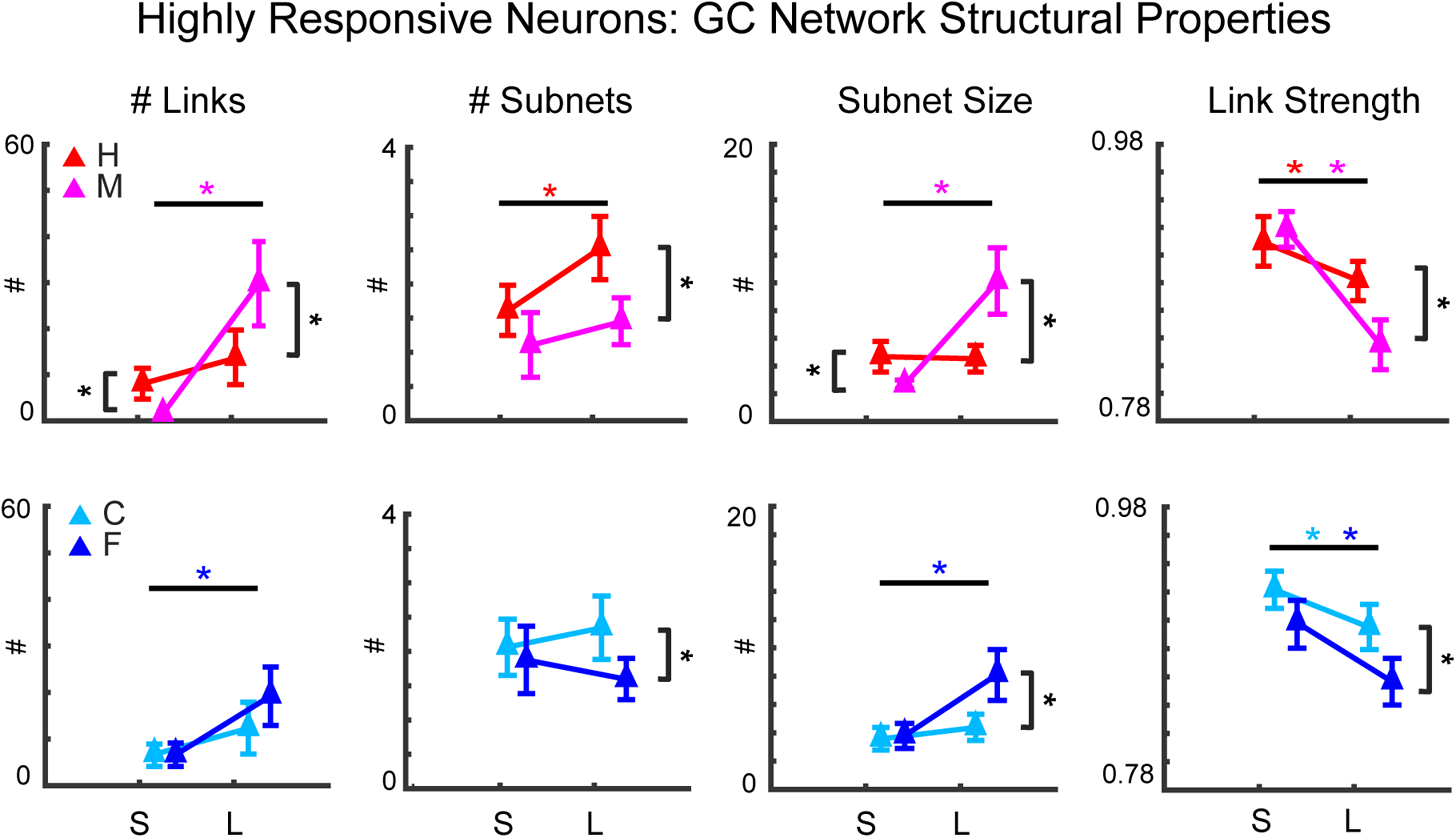
GC network structural properties of highly responsive neurons. Selection of highly responsive neurons is independent of their information content, so the properties of these networks are used as a baseline for comparison (Figure 3D). Comparing S v. L timescales, correct decision (H/C) networks had robust number of links, number of subnetworks, and subnetwork sizes; incorrect decision (M/F) networks had greater number of links and subnetwork size. Both correct and incorrect decision networks had weaker links in L-timescale networks. S-timescale H/C networks had at least as many links as M/F networks, while L-timescale H/C networks had fewer, stronger links. L-timescale H/C networks had smaller, more numerous subnetworks than M/F networks. See also Supplementary Table S1. Panels show means ± 2 SEM. Asterisks indicate statistically significant differences evaluated using Wilcoxon’s signed rank test (p<0.05).

**Figure 3 – Figure Supplement 2.**
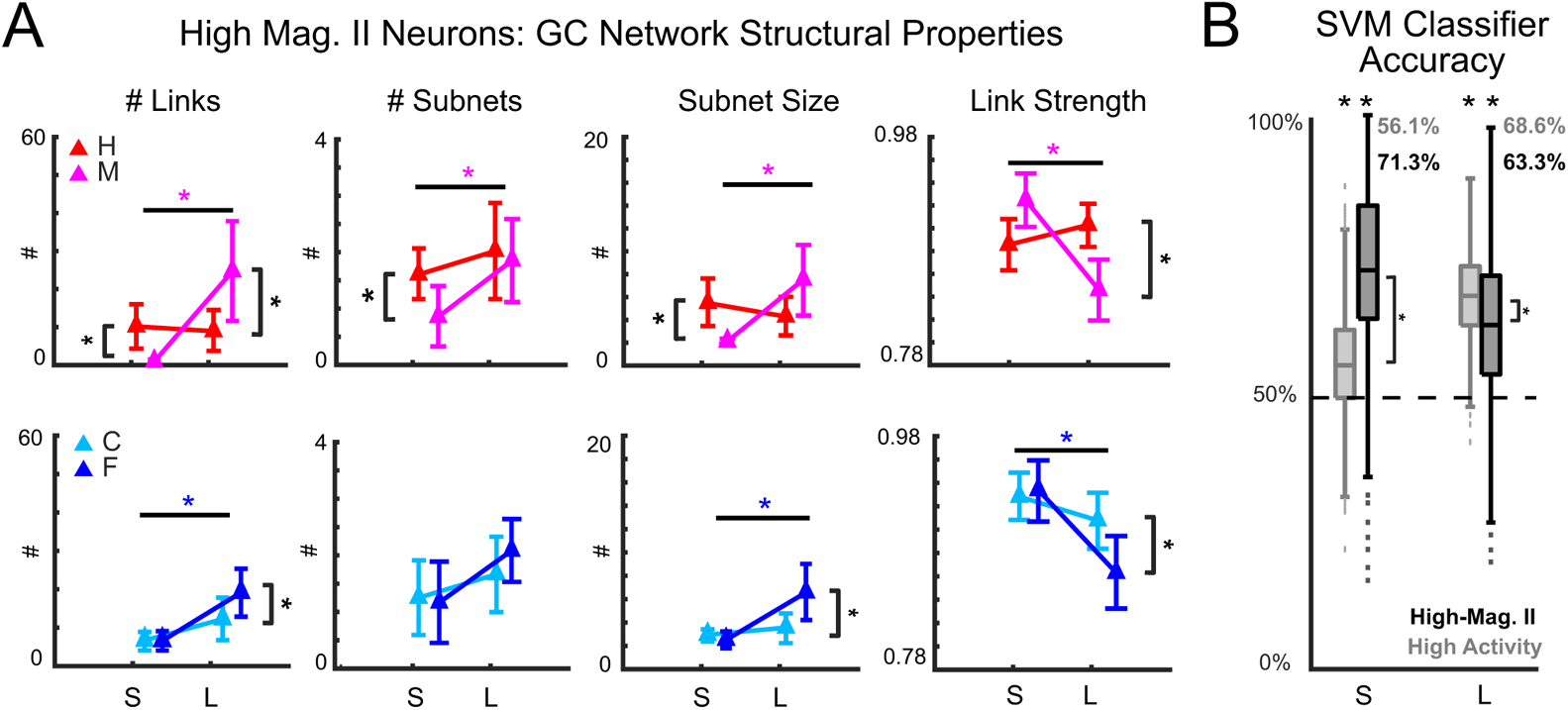
Network structure of neurons with greatest peak *II* magnitudes encode behavioral choice. **A.** Network statistics of greatest peak *II* magnitude neurons differed by timescale and behavioral choice similarly to network statistics of low *II*-peak latency neurons (Figure 3C). See also Supplementary Table S2. **B.** Network statistics were used to train an SVM to classify into correct or incorrect decisions. Across timescale and selection of neurons, decisions were predicted significantly better than chance (p<0.001). S-timescale network structure of high-magnitude *II* neurons was better decoded than of highly responsive neurons (p<0.001). L-timescale network structures had high decoding accuracy, but highly responsive neuronal networks were better decoded (p<0.001). Thus, networks of neurons with greatest peak *II* magnitudes also encode behavioral choice. Panel **A** shows means ± 2 SEM. Asterisks indicate statistically significant differences. Wilcoxon’s signed rank test (p<0.05) was used in **A**. A two-sample t-test (p<0.05) in **B** to compare neurons with high-magnitude *II*, and a one-sample t-test (p<0.05) to compare performance with chance decoding accuracy.

**Figure 3 – Figure Supplement 3.**
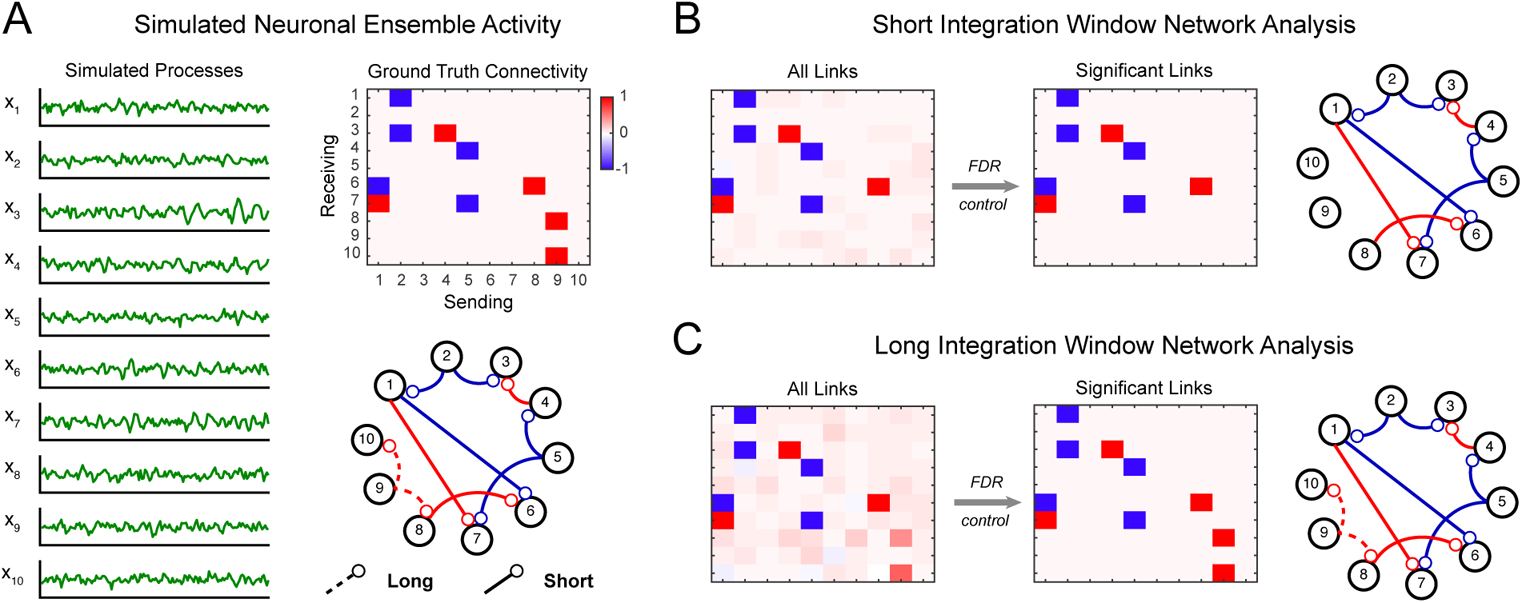
Simulated example for assessing the use of the proposed parametrization of the integration window lengths in Granger causality (GC) network inference. **A.** Simulated responses of 10 neurons, shown averaged over 10 trials of 150 time samples in the left panel, were generated based on an underlying network of long (L) and short (S) timescale interactions (right subpanels). **B.** GC analysis using the short integration window identifies true S-timescale interactions, while expectedly discarding the L-timescale influences. False discovery rate (FDR) control prunes weak spurious interactions and retains significant links. **C.** Employing the L integration window for GC analysis captures both S and L influences, and after FDR control, the true functional connectivity is inferred correctly.

**Figure 6 – Figure Supplement 1.**
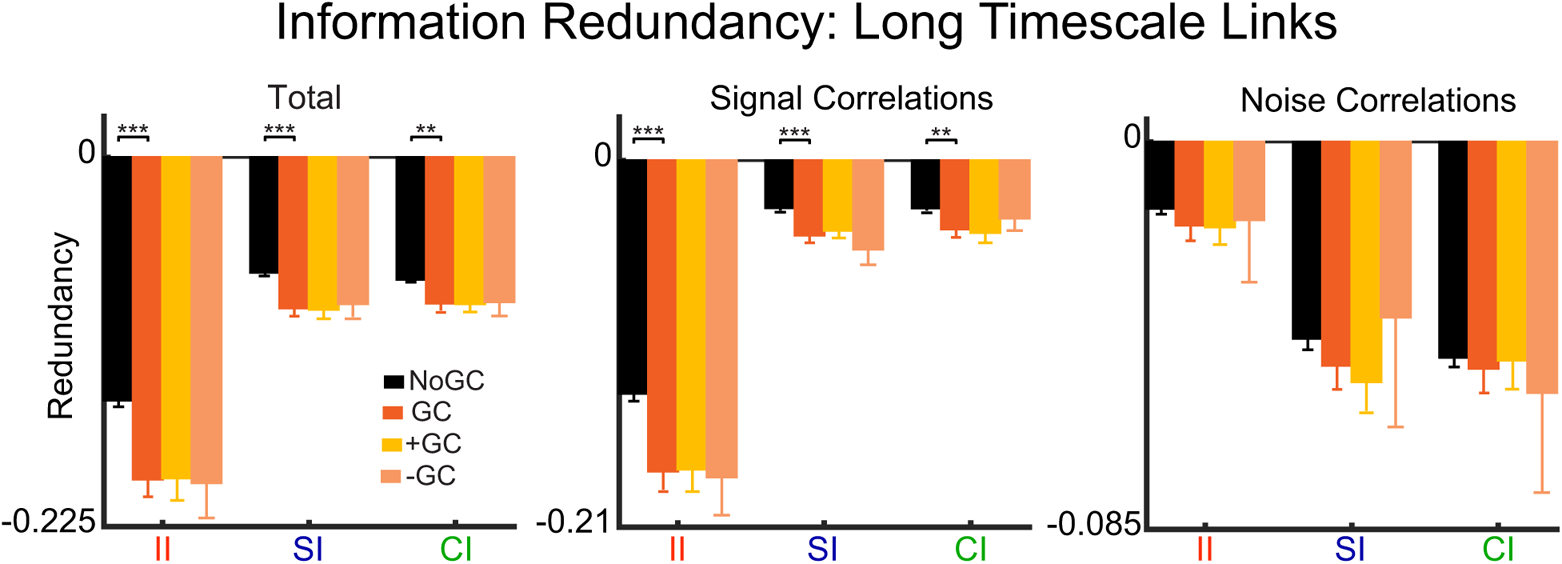
The normalized time-lagged redundancy index showed that GC-linked pairs of neurons in L-timescale networks shared more redundant information (*II, SI, CI*) than GC-unlinked pairs of neurons. The part of redundant information related to signal correlations was more negative for GC-linked pairs of neurons than GC-unlinked neurons (*II, SI, CI*), while noise correlations did not show significant differences in redundant *SI, CI*, or *II* (see Figure 6 in the main text for a comparison to S-timescale networks). We report no difference in redundancy index between groups of positive and negative GC-linked neurons. Statistical comparisons between groups are made with a two-sample t-test (*p<0.05; **p<0.01; ***p<0.001).

## SUPPLEMENTARY TABLES

**Supplementary Table S1.** Highly responsive neurons: statistical comparisons of GC network structure across short (S) and long (L) timescales, and behavioral choice categories — hit (H), miss (M), correct-rejection (C), and false-alarm (F) – using Wilcoxon’s signed rank test (p<0.05). See also Figure 3 Supplement 1.

|  | # Links |  |  | # Subnets |  |  | Subnet Size |  |  | Link Strength |  |  |
| --- | --- | --- | --- | --- | --- | --- | --- | --- | --- | --- | --- | --- |
|  | S | L | p-value | S | L | p-value | S | L | p-value | S | L | p-value |
| H | 8.41 ± 1.605 | 13.91±2.885 | 0.102 | 1.63 ± 0.178 | 2.53 ± 0.229 | **0.003** | 4.64 ± 0.556 | 4.48 ± 0.486 | 0.836 | 0.91 ± 0.009 | 0.88 ± 0.008 | **0.023** |
| M | 1.88 ± 0.453 | 29.91±4.544 | **<0.001** | 1.13 ± 0.233 | 1.47 ± 0.168 | 0.235 | 2.58 ± 0.171 | 10.11±1.174 | **<0.001** | 0.92 ± 0.007 | 0.84 ± 0.009 | **<0.001** |
| p-value | **<0.001** | **0.004** |  | 0.093 | **<0.001** |  | **<0.001** | **<0.001** |  | 0.457 | **<0.001** |  |
| C | 6.72 ± 1.112 | 12.31±2.817 | 0.072 | 2.06 ± 0.206 | 2.34 ± 0.236 | 0.372 | 3.67 ± 0.390 | 4.48 ± 0.467 | 0.183 | 0.92 ± 0.007 | 0.90 ± 0.008 | **0.013** |
| F | 6.63 ± 1.261 | 19.25±3.057 | **<0.001** | 1.88 ± 0.245 | 1.59 ± 0.148 | 0.33 | 3.90 ± 0.454 | 8.23 ± 0.922 | **<0.001** | 0.90 ± 0.009 | 0.86 ± 0.009 | **0.002** |
| p-value | 0.956 | 0.1 |  | 0.56 | **0.01** |  | 0.697 | **<0.001** |  | 0.051 | **0.001** |  |

**Supplementary Table S2.**
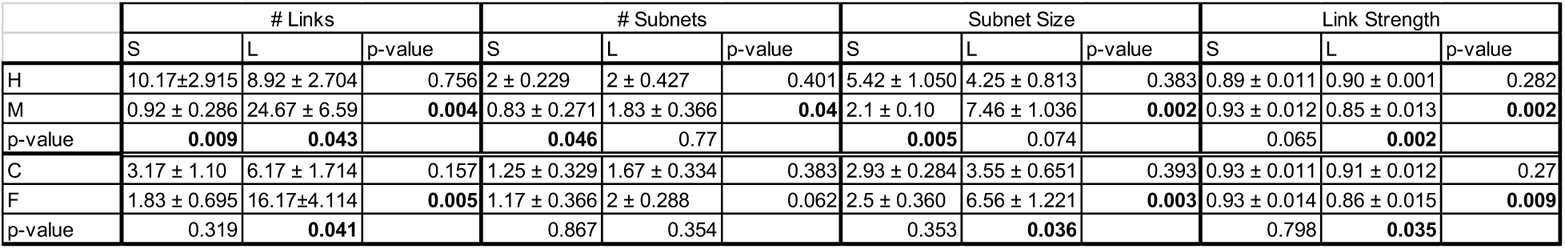
Neurons with greatest *II* peak magnitudes: statistical comparisons of GC network structure across short (S) and long (L) timescales, and behavioral choice categories — hit (H), miss (M), correct-rejection (C), and false-alarm (F) – using Wilcoxon’s signed rank test (p<0.05). See also Figure 3 Supplement 2A.

**Supplementary Table S3.** Supplementary statistical comparisons of correct/incorrect decision GC network structure by stimulus — i.e., hit (H) vs. correct-rejection (C), and miss (M) vs. false-alarm (F) – for short (S) and long (L) timescales using Wilcoxon’s signed rank test (p<0.05). Comparisons performed for networks of low *II*-peak latency neurons (see also Figure 3 and Table 1); highly responsive neurons (see also Figure 3 Supplement 1 and Table S1); *SI/CI* neurons (see also Figure 4 and Table 2); and high-magnitude *II* neurons (see also Figure 3 Supplement 2 and Table S2).

|  | # Links |  | # Subnets |  | Subnet Size |  | Link Strength |  |
| --- | --- | --- | --- | --- | --- | --- | --- | --- |
|  | S | L | S | L | S | L | S | L |
| Low-Latency II Networks |  |  |  |  |  |  |  |  |
| H vs. C | **0.007** | 0.06 | 0.098 | 0.72 | **0.025** | 0.128 | **0.0143** | **0.017** |
| M vs. F | 0.85 | 0.901 | 0.155 | 1 | 0.508 | 0.429 | 0.803 | 0.769 |
| Highly Responsive Networks |  |  |  |  |  |  |  |  |
| H vs. C | 0.391 | 0.694 | 0.113 | 0.57 | 0.157 | 0.998 | 0.277 | 0.229 |
| M vs. F | **0.001** | 0.055 | **0.03** | 0.579 | **0.008** | 0.213 | 0.125 | 0.091 |
| SI/CI Networks |  |  |  |  |  |  |  |  |
| H vs. C | 0.291 | 0.159 | 0.388 | 0.486 | 0.344 | 0.161 | 0.103 | 0.473 |
| M vs. F | 0.207 | 0.767 | 0.758 | 0.534 | 0.17 | 0.484 | 0.649 | 0.346 |
| High Magnitude II Networks |  |  |  |  |  |  |  |  |
| H vs. C | **0.041** | 0.401 | 0.415 | 0.545 | **0.033** | 0.505 | **0.033** | 0.868 |
| M vs. F | 0.242 | 0.288 | 0.472 | 0.597 | 0.301 | 0.651 | 0.771 | 0.464 |

